## Supplementary material for "Traitor-virus-guided discovery of novel antiviral factors": Table S1

**Supplementary Table 1. List of gRNAs targeting 511 candidate restriction factors.**

| sgRNA | Sequence |
|---|---|
| PARP9-gRNA-1 | GTTCATTGTAAGCTGCTGCTC |
| PARP9-gRNA-2 | GCAGCTTACAATGAAAAATC |
| PARP9-gRNA-3 | GAGCTGACACTTTACCATATC |
| APOBEC3G-gRNA-1 | GAGCACAGCCAAGACCTGAG |
| APOBEC3G-gRNA-2 | GTATATTACTGCACATCATGC |
| APOBEC3G-gRNA-3 | GAAGGCTCAAATAGCTCTCTT |
| IFI16-gRNA-1 | GACCAGCCCTATCAAGAAAG |
| IFI16-gRNA-2 | GAAAACGCCCAGTGATAGTGA |
| IFI16-gRNA-3 | GTACACAGAACCCGAAAACAG |
| APOBEC3B-gRNA-1 | GAAAGACCCCTGTGTCCCAA |
| APOBEC3B-gRNA-2 | GAAAACTTTGTGTACAATGA |
| APOBEC3B-gRNA-3 | GCCAGACCTACTTGTGCTATG |
| TRIM5-gRNA-1 | GTTTCGAGCTCCTGATCTGAA |
| TRIM5-gRNA-2 | GTCTAGCATTCCTTTCAGATC |
| TRIM5-gRNA-3 | GTTTCTCACAGGTGAAGCTCC |
| OAS1-gRNA-1 | GAGGCCGATCTGACGCTGACC |
| OAS1-gRNA-2 | GAATCTATGTCAAGCTCATCG |
| OAS1-gRNA-3 | GTCAGTACGAAGCTGAGCGCA |
| IFI44-gRNA-1 | GTGAATTCTGTCCTTCAGCGA |
| IFI44-gRNA-2 | GAGAAAGAAGGCGGCCTGTGC |
| IFI44-gRNA-3 | GCCTGATGCGTTACATGCCCT |
| EIF2AK2-gRNA-1 | GAGCTGTTGAGATACTTAATA |
| EIF2AK2-gRNA-2 | GCAACCTACCTCCTATCATG |
| EIF2AK2-gRNA-3 | GAAGTATCTCAACAGCTAATT |
| APOL1-gRNA-1 | GCTGCCCTCACTCCCACACCA |
| APOL1-gRNA-2 | GCCAAGCTCACCAGATGCAG |
| APOL1-gRNA-3 | GATCCTCAAAGTAAGCCCCT |
| MKI67-gRNA-1 | GAAAATGAAAGTCTTCAGAA |
| MKI67-gRNA-2 | GTCTAGCTTCTCTTCTGACCC |
| MKI67-gRNA-3 | GATCAAAAGGAGCGGGGTCGA |
| SP100-gRNA-1 | GCTCCAACTAAGTCTTGAACA |
| SP100-gRNA-2 | GCAGCCTGTCATCTACACCC |
| SP100-gRNA-3 | GACATTTAACCTGCCAGTTC |
| PARP14-gRNA-1 | GTGGTTCTTTAACATCTTCTT |
| PARP14-gRNA-2 | GAAAGAAGATGTTAAAGAACC |
| PARP14-gRNA-3 | GCTGAAGTTGTTTAATTGAA |
| LILRA1-gRNA-1 | GCCCTTCTTCACAATCTCCTG |
| LILRA1-gRNA-2 | GAGCTGACATCAGAAACACAC |
| LILRA1-gRNA-3 | GGGCGGTATCGCTGTTTCTA |
| HERC6-gRNA-1 | GAAAATAATACAAGTTTCCTG |
| HERC6-gRNA-2 | GCACTCCCTGGCATTATCAAA |
| HERC6-gRNA-3 | GTGCACTGAAGAATCTAGGTG |

| | |
|---|---|
| GZMA-gRNA-1 | GCGTGTGGCTGGGTCATAGCA |
| GZMA-gRNA-2 | GTTCCGAGGGGTCACTTCCTT |
| GZMA-gRNA-3 | GAATAAATGCGGAGACCCTCG |
| IGFL2-gRNA-1 | GTACTCACCGATGACTTCCCT |
| IGFL2-gRNA-2 | GAGACAGACTGACACATAAGC |
| IGFL2-gRNA-3 | GTTACTTACCCAGCACACTCC |
| OAS2-gRNA-1 | GTGATCCTCTTGATAAAGCAC |
| OAS2-gRNA-2 | GATGCCAGTGCTTTATCAAG |
| OAS2-gRNA-3 | GACCCAACCAATAATGTGAG |
| LAIR1-gRNA-1 | GTTCTCTTCCAGTGCTCTGCC |
| LAIR1-gRNA-2 | GCTGTTTACAGAAACCTCTGG |
| LAIR1-gRNA-3 | GCCCTCCCAGGACCCACGCAG |
| RNASE3-gRNA-1 | GTTAATTGCCCGCATTGCAA |
| RNASE3-gRNA-2 | GACTGGAACCACAGGATACCG |
| RNASE3-gRNA-3 | GCACTGAGCCCTCGTAAACTG |
| LILRB2-gRNA-1 | GTTTGCCCTGACGTCGATGT |
| LILRB2-gRNA-2 | GTCAAGGTCTGGGAAGGCACC |
| LILRB2-gRNA-3 | GACCGGTCCCATCTCCACACC |
| APOL3-gRNA-1 | GATATACCCTGGAACCCAAA |
| APOL3-gRNA-2 | GCGCAGTCACGAATCTCTTCC |
| APOL3-gRNA-3 | GACTCTCTCCCGGAAGTATT |
| TNFSF10-gRNA-1 | GTACAGACTCCAAGAATGAAA |
| TNFSF10-gRNA-2 | GATGTGAGCTGCTACTCTCTG |
| TNFSF10-gRNA-3 | GACTACCTTGAACTGTAGAAA |
| C2orf16-gRNA-1 | GCCGACGCAGTCCCCTTAAGG |
| C2orf16-gRNA-2 | GTAAAGTCTGTGACGATACCA |
| C2orf16-gRNA-3 | GCGCCTGATGTAGACTTGTAC |
| IFI44L-gRNA-1 | GATCATCCACTTTAGTAAGCA |
| IFI44L-gRNA-2 | GAATCTTACCTGATATCTGTC |
| IFI44L-gRNA-3 | GTTATCCTCTTTATGTCGTCT |
| CASP10-gRNA-1 | GTAGACCTCCCTAAGTTTCC |
| CASP10-gRNA-2 | GTGAAGATAATCTGACATGCC |
| CASP10-gRNA-3 | GACTGCTGCCCACCCGACAA |
| SIGLEC9-gRNA-1 | GTTGACCATGACTGTCTTCCA |
| SIGLEC9-gRNA-2 | GTCCTACCTGTGCCGTCTCCT |
| SIGLEC9-gRNA-3 | GGACACGGAGGTCCCTATCC |
| RARRES3-gRNA-1 | GGCCACACCAACTTCAACCT |
| RARRES3-gRNA-2 | GTTGGTGTGGCCACGGCGCT |
| RARRES3-gRNA-3 | GAGATGGCTACGTGATCCATC |
| MLKL-gRNA-1 | GAAGAACCCTACCATCCACAG |
| MLKL-gRNA-2 | GATCCCGCAAGAGCAAATCA |
| MLKL-gRNA-3 | GTTCTGAGAAGATCCGCAAGC |
| CASP5-gRNA-1 | GACGAAAAGAATCTCACAGCC |
| CASP5-gRNA-2 | GTAGTGAGCCCCATTCCTTGC |

| | |
|---|---|
| CASP5-gRNA-3 | GTTTGTAGATTCTGCTGACTC |
| TLR4-gRNA-1 | GACCTGAGCTTTAATCCCCTG |
| TLR4-gRNA-2 | GCCCCTTCTCAACCAAGAACC |
| TLR4-gRNA-3 | GCCAGCTTTCTGGTCTCACGC |
| FCGR3A-gRNA-1 | GAAAAAGCCCCCTGCAGAAGT |
| FCGR3A-gRNA-2 | GAAAGCCACACTCAAAGACAG |
| FCGR3A-gRNA-3 | GATCTCATCATTCTTTCCACC |
| SP110-gRNA-1 | GTCTTCTTCCGCATTCATTT |
| SP110-gRNA-2 | GTACCTTGCACAGTGCTAGTG |
| SP110-gRNA-3 | GAGAATGTTGTGCACCACTC |
| DEFB1-gRNA-1 | GCCGATCTTTACCAAAATTCA |
| DEFB1-gRNA-2 | GACAGGTGGTAACTTTCTCAC |
| DEFB1-gRNA-3 | GTAACAGGTGCCTTGAATTT |
| CASP1-gRNA-1 | GCTTGAGAGTCTTGCATATTA |
| CASP1-gRNA-2 | GATGGAAACAAAAGTCGGCAG |
| CASP1-gRNA-3 | GAAAGCTGTTTATCCGTTCCA |
| CD3E-gRNA-1 | GAGATAAAAGTTCGCATCTTC |
| CD3E-gRNA-2 | GTATTATGTCTGCTACCCCAG |
| CD3E-gRNA-3 | GAGGGCATGTCAATATTACTG |
| MS4A4A-gRNA-1 | GAATTCCTGCTGCAATTGACA |
| MS4A4A-gRNA-2 | GCAACTTACCCCAAGGACTT |
| MS4A4A-gRNA-3 | GAATTGTGTACCCGATATACA |
| APOBEC3F-gRNA-1 | GTGGCCTCCCCAGCTGACCGC |
| APOBEC3F-gRNA-2 | GCCCACCACATGGGACAGCGC |
| APOBEC3F-gRNA-3 | GAAAAACCTACGCAAAGCCTA |
| C1orf162-gRNA-1 | GCTAGTTGTTGGGGCTGCTG |
| C1orf162-gRNA-2 | GCTGCAGCTTTCATCCATCCC |
| C1orf162-gRNA-3 | GCAGTAGAACCCCAGCACAAA |
| BRCA2-gRNA-1 | GAAATCAAGAAAAATCCTTAA |
| BRCA2-gRNA-2 | GACTTATTAAATGAATTTGAC |
| BRCA2-gRNA-3 | GTCTTACCGAAAGGGTACAC |
| RNASE2-gRNA-1 | GACTGGAACCACCGGATACTG |
| RNASE2-gRNA-2 | GGGAGGTCATATTGATGTGC |
| RNASE2-gRNA-3 | GTTAATGACCTGCATTGCAT |
| TAPBP-gRNA-1 | GTGTCCCACTTCTACCCTTC |
| TAPBP-gRNA-2 | GAAGCGGCTCATCTCGCAGTG |
| TAPBP-gRNA-3 | GCCCCGGGGATACCGCCTGA |
| GGH-gRNA-1 | GACGCTCAGATTATGCTAAAG |
| GGH-gRNA-2 | GTGACTGCCAATTTCCATAAG |
| GGH-gRNA-3 | GTGCGTCCTATGTAAAGTACT |
| RIPK3-gRNA-1 | GCATGGAGAACGGCTCCTTGT |
| RIPK3-gRNA-2 | GAACTGTTTGTTAACGTAAAC |
| RIPK3-gRNA-3 | GTCCTTACCTGTAGACGTCAC |
| SSX7-gRNA-1 | GAATACTTCTCTAAGAAAGAG |

| | |
|---|---|
| SSX7-gRNA-2 | GTTCTGTGAGCCAGATGCTTC |
| SSX7-gRNA-3 | GAGCCTGCAAAAAGTCATCTG |
| HLA-B-gRNA-1 | GCTCCGATGACCACAACTGCT |
| HLA-B-gRNA-2 | GCGTACTGGTCATGCCCGCGG |
| HLA-B-gRNA-3 | GCCTGGCTGTCCTAGCAGTTG |
| SP140-gRNA-1 | GGCCAAGCTCATGGCCCAGC |
| SP140-gRNA-2 | GCACTTGCCATCTGCCCCTGC |
| SP140-gRNA-3 | GTATTGTGTACTCAGTGAAC |
| NKG7-gRNA-1 | GTTGCTTTGAGCACCGATTTC |
| NKG7-gRNA-2 | GCTGCGGTGGTTGAGACAAGC |
| NKG7-gRNA-3 | GTGGCGGTGTACACCAGCGAG |
| HLA-A-gRNA-1 | GCCACCCCATCTCTGACCATG |
| HLA-A-gRNA-2 | GTCCCTCCTTACCCCATCTCA |
| HLA-A-gRNA-3 | GCCAGTCACAGACTGACCGAG |
| HAVCR1-gRNA-1 | GCATGAAAATCACCGTATCAT |
| HAVCR1-gRNA-2 | GAACCCACCCTACGACACTGC |
| HAVCR1-gRNA-3 | GCCATTGTACTCTTACACAAC |
| LILRA6-gRNA-1 | GCCCTCCCTCCTGACCCTGC |
| LILRA6-gRNA-2 | GCAGACTCACCTGCCTGCACG |
| LILRA6-gRNA-3 | GCTCTCTTCCAGGGCTGAGTC |
| COA1-gRNA-1 | GCAGTTGAAGATTCCTGTCTC |
| COA1-gRNA-2 | GTGTCGATGAGCTTGAGATAA |
| COA1-gRNA-3 | GATTATCTCAAGCTCATCGAC |
| TCHHL1-gRNA-1 | GATGTTTGACACTCAAGAACC |
| TCHHL1-gRNA-2 | GACTCATGACCAACCAGTTG |
| TCHHL1-gRNA-3 | GTGCCCCGTTACTGTCCTCAC |
| FCRL3-gRNA-1 | GCAGCAGCAGCCACAGAAGCA |
| FCRL3-gRNA-2 | GTTACCTGATTGTTCTCTTCC |
| FCRL3-gRNA-3 | GCTCCTGGAAGAGAACAATC |
| KRTAP19-2-gRNA-1 | GGCCACAGAGGCTGTGGAGA |
| KRTAP19-2-gRNA-2 | GAGCCATCTCCACAGCCTCTG |
| KRTAP19-2-gRNA-3 | GGTGGCATGTGCTATGGCTA |
| MT1H-gRNA-1 | GCTGCTGCTCCTGTTGCCCCC |
| MT1H-gRNA-2 | GCCCCTTTGCAGATGCAGCCC |
| MT1H-gRNA-3 | GCCCGCACTCACTCTTCTTGC |
| NDUFB6-gRNA-1 | GTTATTACAGGAAAAACCATA |
| NDUFB6-gRNA-2 | GAAGACTTACAGGGAATATTC |
| NDUFB6-gRNA-3 | GTTCCTAGGGTGATACAATTC |
| MARCO-gRNA-1 | GTTTGGTGAAAAGCAGCTTGT |
| MARCO-gRNA-2 | GGTGAACTTCTCCCTAGCTG |
| MARCO-gRNA-3 | GTGACGCGGACCCAGGTGAGT |
| GBP5-gRNA-1 | GTACTGTGAACAAAATTGATC |
| GBP5-gRNA-2 | GTGATATCCAGATCTTTGCAC |
| GBP5-gRNA-3 | GGTGTGTGCCTCATCCCAAC |

| | |
|---|---|
| TMEM126B-gRNA-1 | GTCTATGATTTCTATGACCAT |
| TMEM126B-gRNA-2 | GAAGAACTTACTTGGTTGCC |
| TMEM126B-gRNA-3 | GCCTTGAAGCAGCGTCTGAAC |
| DTX3L-gRNA-1 | GTGTTGATGCATATCACCAGA |
| DTX3L-gRNA-2 | GAGTCTCAGATGTCATCACT |
| DTX3L-gRNA-3 | GTATGCAGTTCGCTGTATTCC |
| CD4-gRNA-1 | GCCTGCTGGAATCCAACATCA |
| CD4-gRNA-2 | GAGCTCCAGGATAGTGGCACC |
| CD4-gRNA-3 | GTGGGTCCTTACCCTTGATGT |
| PSG11-gRNA-1 | GTCAAATAATTATATATGGAC |
| PSG11-gRNA-2 | GCCAGCCACCCAAAGTGTCCG |
| PSG11-gRNA-3 | GAGCTGCATCCTATGAGTCAT |
| PRG3-gRNA-1 | GGCTTAGAAGGAGCAGACGA |
| PRG3-gRNA-2 | GCCCCCTCACCTTTGGTGCAT |
| PRG3-gRNA-3 | GTCTGGATTGGAGGCAACCTC |
| APOBEC3A-gRNA-1 | GCTTACCATCGTAGGTCATGA |
| APOBEC3A-gRNA-2 | GTTTGCAGTGCCTCCTTATAT |
| APOBEC3A-gRNA-3 | GATGGAAGCCAGCCCAGCATC |
| ANKRD20A3-gRNA-1 | GTACCGTCTCAGCCTATCAA |
| ANKRD20A3-gRNA-2 | GAATTTGCAAGAAAGAGAAAA |
| ANKRD20A3-gRNA-3 | GTGCTTCAATATGTGCACCA |
| APOL6-gRNA-1 | GCAACGTACCACTCACGATGC |
| APOL6-gRNA-2 | GCGCCAGTGTTGTTCCCGCAA |
| APOL6-gRNA-3 | GCCTCACTTTCTCTCTCCGCC |
| SIGLEC7-gRNA-1 | GGTCCTGTTCGTGGTCACGC |
| SIGLEC7-gRNA-2 | GGCAAAATGAGGCCTGTATC |
| SIGLEC7-gRNA-3 | GCACCTGTGTACTCCTGTTGC |
| CENPF-gRNA-1 | GACAGCTTGACAAACTGAAGA |
| CENPF-gRNA-2 | GACACCAAGTCAATATTATAG |
| CENPF-gRNA-3 | GGTTGAAAATGAAAAAACCG |
| C6orf1-gRNA-1 | GAGAGTGGCTACCAGCACAAG |
| C6orf1-gRNA-2 | GTGTATGTACAGAAGAAAAAG |
| C6orf1-gRNA-3 | GTACAGCTATGACCCAGCTG |
| PTPRC-gRNA-1 | GTAATTCTTACCAGTGGGGGA |
| PTPRC-gRNA-2 | GAGCCCAACACCTTCCCCCAC |
| PTPRC-gRNA-3 | GACTCCATCTAAGCCAACATG |
| GZMH-gRNA-1 | GTTCCTTTCAGAGGAGATCAT |
| GZMH-gRNA-2 | GTGGCGGCATCCTAGTGAGAA |
| GZMH-gRNA-3 | GCATGGAAGAGACGTTCACAC |
| CD58-gRNA-1 | GATGTTAAGTTGTAGATAGTG |
| CD58-gRNA-2 | GCATGTTGTAATTACTGCTAA |
| CD58-gRNA-3 | GCTAACTTGTGCATTGACTAA |
| CCR3-gRNA-1 | GAGTTATGCCCCCTGACATAG |
| CCR3-gRNA-2 | GCACGTCATCATAGTAGGATG |

| | |
|---|---|
| CCR3-gRNA-3 | GCTGTTAAGAGCACACAAGCC |
| POLQ-gRNA-1 | GATGCTGGGAGACTCTCACCG |
| POLQ-gRNA-2 | GCCGAGTAATATAGCAAATCT |
| POLQ-gRNA-3 | GAATAAAAGTAGACGGTTATA |
| CLEC12A-gRNA-1 | GCAATACATGTTATTTAAGTC |
| CLEC12A-gRNA-2 | GTTATGCCAAATCCATCTCCT |
| CLEC12A-gRNA-3 | GCTCCAGCTCCCTCTCATGTA |
| SGOL2-gRNA-1 | GAGTGATGGAGTGCCCAGTGA |
| SGOL2-gRNA-2 | GCAAATGTCTCTTAATTCCTG |
| SGOL2-gRNA-3 | GAGTGAGCCAGTTTCCATCAC |
| BTN3A2-gRNA-1 | GCTGAGCAAGGAGTGAGCAGC |
| BTN3A2-gRNA-2 | GCTTTCATGTCTCCCTCCTCT |
| BTN3A2-gRNA-3 | GTAACTTACTGAGTTCCTCC |
| LAIR2-gRNA-1 | GTTCTCTTCCAGTGCTCTGCC |
| LAIR2-gRNA-2 | GATGACTTACCCTCCTGCGTG |
| LAIR2-gRNA-3 | GCATTCATGGTGCATCAAATC |
| TROAP-gRNA-1 | GAGGGTCAGCCACAAACTCTA |
| TROAP-gRNA-2 | GTCCAGGGACCATAGAGTTTG |
| TROAP-gRNA-3 | GGGCGTTGAATATTGAGCGG |
| KRTAP5-4-gRNA-1 | GCTGTAGAACGATGCCATATC |
| KRTAP5-4-gRNA-2 | GTAAGCGTCCCACCTGCAACA |
| KRTAP5-4-gRNA-3 | GTGAGCCCCAAATCATTGCTC |
| PTPRJ-gRNA-1 | GCGGCTGCCTCCGCGCTCGCC |
| PTPRJ-gRNA-2 | GCAGCCTTCCTCACACCCGTG |
| PTPRJ-gRNA-3 | GCGCGCGGGGCATGAAGCCGG |
| RAET1L-gRNA-1 | GTCCACCACCTCTCTCAGTAC |
| RAET1L-gRNA-2 | GTCATCCCTAAGTTCAGACC |
| RAET1L-gRNA-3 | GTCTTCTCTGAGTCAAAGAGT |
| APOBEC3C-gRNA-1 | GAATCCCTTCCAAGGCTTGAA |
| APOBEC3C-gRNA-2 | GCTACGGGAGAGTCTCCAGTG |
| APOBEC3C-gRNA-3 | GTCTCCTAACACAAAGTACC |
| ZBP1-gRNA-1 | GAGTCACTTTCCTCGAGACAT |
| ZBP1-gRNA-2 | GCTCGAAGCTCACCCCCAAGC |
| ZBP1-gRNA-3 | GATCAGTCCCGCCCAAGCACC |
| AIM2-gRNA-1 | GTTTGACCTAAGTGACAACAC |
| AIM2-gRNA-2 | GAAAAGTCTCTCCTCATGTTA |
| AIM2-gRNA-3 | GCCTTATCCTACCTTAACATG |
| IFNAR1-gRNA-1 | GCTGGAGCCACTGAACTTGAA |
| IFNAR1-gRNA-2 | GTAGATGACAACTTTATCCTG |
| IFNAR1-gRNA-3 | GATCTAATGTTAAAGACTGG |
| C4BPA-gRNA-1 | GCTTTACTTACTTTCACATTG |
| C4BPA-gRNA-2 | GTCCAGCCAACTCCTCTATCT |
| C4BPA-gRNA-3 | GACAAACGATGCAGACACCC |
| FCGR1B-gRNA-1 | GTCTGTGGATTTCCAGCTGTA |

| | |
|---|---|
| FCGR1B-gRNA-2 | GTCTCCAAGTAGACACCACAA |
| FCGR1B-gRNA-3 | GTCACCCACTTGCCCATCAAC |
| CEACAM1-gRNA-1 | GTCTGATTGTTTATCCACCAC |
| CEACAM1-gRNA-2 | GTCACTGCGGTTCGCACTCAC |
| CEACAM1-gRNA-3 | GTGTCTCTCGACCGCTGTTTG |
| CD72-gRNA-1 | GTTCAAGTGCCCGCAGTCCT |
| CD72-gRNA-2 | GTGATGCATTATCCATCCCGA |
| CD72-gRNA-3 | GGTATCGCAGGCAGGTTGTG |
| PRG4-gRNA-1 | GAAACAGACAGCAGCAACAAC |
| PRG4-gRNA-2 | GATTCAGCAAGTTTCATCTCA |
| PRG4-gRNA-3 | GTGCTTTGAGTCCTTCGAGAG |
| CCNB3-gRNA-1 | GACTGGCCAAAAAGAATAAG |
| CCNB3-gRNA-2 | GTTGTTTCCAAGAAGATAAAC |
| CCNB3-gRNA-3 | GTCATCATGACCCATCTGAAA |
| CEACAM3-gRNA-1 | GTGTGAGCAGAAGCCCCTGCC |
| CEACAM3-gRNA-2 | GCTCCCACAGCTCTGCCTTCT |
| CEACAM3-gRNA-3 | GATGCATTCTCTGTGGGGAG |
| RPS21-gRNA-1 | GAGATGATTCCATTCTCCGAT |
| RPS21-gRNA-2 | GTTCTAGGTTGACAAGGTCAC |
| RPS21-gRNA-3 | GACCGATGATGCGATTGCTAG |
| SPAG5-gRNA-1 | GTCATAATACTCATTTCTGCC |
| SPAG5-gRNA-2 | GCTTCTTTAGACTCTTGTAGC |
| SPAG5-gRNA-3 | GTTTCACCCGAGTAGCATCAA |
| CD38-gRNA-1 | GTCTGGCCCATCAGTTCACAC |
| CD38-gRNA-2 | GCCAGCGGGACATGTTCACCC |
| CD38-gRNA-3 | GTGAAAGCATCCCATACACTT |
| TPRX1-gRNA-1 | GCGCCGCTGGAAGGATTCCCG |
| TPRX1-gRNA-2 | GCCATTCCGTGGCCCAATCCC |
| TPRX1-gRNA-3 | GCCTGCAGCTGTTGCTCCCTG |
| LILRA5-gRNA-1 | GTGTGCAGATGGATGAGACCA |
| LILRA5-gRNA-2 | GCGAGGCAGAGCAGAACCATG |
| LILRA5-gRNA-3 | GGAGGCCCAGGAATACCGTC |
| SIRPB1-gRNA-1 | GTCGGATGGTCTCAGACAAGT |
| SIRPB1-gRNA-2 | GAGCCAAACCCTCTGCCCCCG |
| SIRPB1-gRNA-3 | GCCAGCAGTAGCGTCATCAGC |
| NMI-gRNA-1 | GTATAAAAGATGAACAAAATA |
| NMI-gRNA-2 | GTGACACACAACAAATTCTTA |
| NMI-gRNA-3 | GACAACTGGCTGTCATTCTC |
| MNDA-gRNA-1 | GCGCAAGAAACAAACTGACAT |
| MNDA-gRNA-2 | GTACATCAATTAAGTCCTTAC |
| MNDA-gRNA-3 | GTTCTTGCATCCACCTGCCGT |
| TAP1-gRNA-1 | GGGATCAGTGTCCCTCACCA |
| TAP1-gRNA-2 | GAGAGTTAAGTTTCGAGTGA |
| TAP1-gRNA-3 | GCTTGTAGAATCCAGTCAGTG |

| | |
|---|---|
| LILRB4-gRNA-1 | GCAGACTCACCTGCCTGCATG |
| LILRB4-gRNA-2 | GCCCAGGACCCACATGCAGGC |
| LILRB4-gRNA-3 | GCCCTGCATAGTCCTCTGTCA |
| CDKN2A-gRNA-1 | GTTGTAGAAGCAGGCATGCGT |
| CDKN2A-gRNA-2 | GCCCAACGCACCGAATAGTTA |
| CDKN2A-gRNA-3 | GTCTTGGTGACCCTCCGGATT |
| PYHIN1-gRNA-1 | GAGATGTCCAAAGAGCAGACT |
| PYHIN1-gRNA-2 | GACAGGAAGTATGGCTGTAGT |
| PYHIN1-gRNA-3 | GACGACAATCTATGAAATTC |
| IFIT1-gRNA-1 | GTGCGATCTCTGCCTATCGCC |
| IFIT1-gRNA-2 | GCCAATTTGTAGACGAACCCA |
| IFIT1-gRNA-3 | GTTAAGCGGACAGCCTGCCTT |
| APOL4-gRNA-1 | GAGGGTGCAGCAAAACCATCC |
| APOL4-gRNA-2 | GTGCGTGTGGCTGAATTGCCC |
| APOL4-gRNA-3 | GTGCTGACTAGCGATGAAGCC |
| HLA-C-gRNA-1 | GCTTCCTCCTACACATCATAG |
| HLA-C-gRNA-2 | GCTCTCAGCTGCTCCGCCGCA |
| HLA-C-gRNA-3 | GTCACCGCTATGATGTGTAGG |
| MT1X-gRNA-1 | GCTGCTGCTCCTGCTGCCCTG |
| MT1X-gRNA-2 | GTCCCTTTGCAGATGCAGCCC |
| MT1X-gRNA-3 | GCCTGCACTCACTCTTCTTGC |
| LAMP2-gRNA-1 | GACCAGAACGAGCCCTGAGCC |
| LAMP2-gRNA-2 | GCAAGAACATCCCAGTAGTGT |
| LAMP2-gRNA-3 | GATGATGTTGTCCAACACTAC |
| C17orf47-gRNA-1 | GGGAGACTGCAGCACTTCGA |
| C17orf47-gRNA-2 | GTGAAACCAAGCCCTCCGCAA |
| C17orf47-gRNA-3 | GCATCCCCCAAGCTCCCACAG |
| HRC-gRNA-1 | GTGAGCACTGCGACCAGTGCC |
| HRC-gRNA-2 | GCGCAGGTCCACAGGATGCTC |
| HRC-gRNA-3 | GCCATACTCCTGAGCATCCTG |
| FCRL5-gRNA-1 | GTTCAGTTCCGGTGTCTCGCC |
| FCRL5-gRNA-2 | GCTCACCGCCTTACTGCGCTG |
| FCRL5-gRNA-3 | GACAAGTCAGTCCGCTGTGAA |
| ZNF28-gRNA-1 | GTTTCAGTCGCAAATCACACA |
| ZNF28-gRNA-2 | GTCAGGGATGGCTCTTCCTC |
| ZNF28-gRNA-3 | GTCTTTACAGGGATGTGATGC |
| NDUFS5-gRNA-1 | GTCGGCCTTAACATAGATCGA |
| NDUFS5-gRNA-2 | GTCAACCATCGATCTATGTTA |
| NDUFS5-gRNA-3 | GTCCCCCTTGCCAATGTGGTG |
| CENPE-gRNA-1 | GAATGTCATTTATCAAGTTGA |
| CENPE-gRNA-2 | GTATAAGCTTACCAAAATTGA |
| CENPE-gRNA-3 | GATCATCGATTCTGCCATACA |
| SIGLEC10-gRNA-1 | GGTGTTGTGGTCCTGGGGTC |
| SIGLEC10-gRNA-2 | GCACGTGAAATGCTGATAACA |

| | |
|---|---|
| SIGLEC10-gRNA-3 | GCAGCTTCACGCCCAGACCCC |
| XRRA1-gRNA-1 | GTCAGAAAACCTGCTGCCTCT |
| XRRA1-gRNA-2 | GAAAGCTTACCTAGAGGCAGC |
| XRRA1-gRNA-3 | GAGAGTCTCGGCGGGAAAATC |
| CTAG1A-gRNA-1 | GAGCAGCTTTCCCTGTTGATG |
| CTAG1A-gRNA-2 | GGCTTAGCGCCTCTGCCCTG |
| CTAG1A-gRNA-3 | GCATGCCTTTCGCGACACCCA |
| AHSG-gRNA-1 | GTTCCAGCTGTTGAAACTAGA |
| AHSG-gRNA-2 | GCAAAATGTGATTCCAGTCC |
| AHSG-gRNA-3 | GTGAGATTGAAATAGACACCC |
| GIMAP7-gRNA-1 | GACAGCCTTGATCAATGCAA |
| GIMAP7-gRNA-2 | GCAACAGCGAACACCATCCT |
| GIMAP7-gRNA-3 | GCTTCTTGTTGTAGACACTCC |
| GBP4-gRNA-1 | GACCTTCTCAGAGAGCATGAA |
| GBP4-gRNA-2 | GAAAGCCGCTTAAGCTCAGCC |
| GBP4-gRNA-3 | GAAGTAGAGATTGTGTCCTCC |
| PAGE1-gRNA-1 | GAGGCCCAACTCCTGGACATC |
| PAGE1-gRNA-2 | GAACAGGTTCACCCGAAGAC |
| PAGE1-gRNA-3 | GTCATCTGCTCTTCATTTCGC |
| MX2-gRNA-1 | GTTCCGGTAGCTGATCCTTC |
| MX2-gRNA-2 | GTGTGGTGGCACTGTGCCGAA |
| MX2-gRNA-3 | GACAACCAGCCCCGAGACAT |
| ZWINT-gRNA-1 | GCTACCTGTTGTAGCTGCCAC |
| ZWINT-gRNA-2 | GATACCACCACAAACTCAACC |
| ZWINT-gRNA-3 | GGCCTCTACGTGCTCCCTGT |
| LRRC37A-gRNA-1 | GCAGGGACAGCGAAGCCCCAA |
| LRRC37A-gRNA-2 | GTCACTCTCCTCCTCCTCCGT |
| LRRC37A-gRNA-3 | GTTTACCCAGGGTCCCTGTTG |
| PHF11-gRNA-1 | GAATCTGCAGCTGTATTCTTC |
| PHF11-gRNA-2 | GACTTGTGGAATGTGAGGATC |
| PHF11-gRNA-3 | GACGCAGTTCCACAGTCTGA |
| VRK2-gRNA-1 | GTGCAAGACATGTAGTAAAAG |
| VRK2-gRNA-2 | GACATGTCTTGCATCTTTCTC |
| VRK2-gRNA-3 | GAGACCAGATCCATAAAACAG |
| USP26-gRNA-1 | GCATTCAAGCTATAGCGTTTG |
| USP26-gRNA-2 | GTCATGCATCATGAACGCCAC |
| USP26-gRNA-3 | GATGGCTGCCCTATTCCTACG |
| TOPBP1-gRNA-1 | GTTTAAAAACTTCACAAAAAA |
| TOPBP1-gRNA-2 | GAGAGCTTTAAAAAAACATT |
| TOPBP1-gRNA-3 | GTGTCTTTGATCACCTCAAAA |
| TFB2M-gRNA-1 | GACTTGAAGCTGGTGCCAAAG |
| TFB2M-gRNA-2 | GCTTAATACATGATACAAGTC |
| TFB2M-gRNA-3 | GTAGAAACTAATGGCAGATCC |
| ZNF429-gRNA-1 | GACCATTGACATTTACAGATG |

| | |
|---|---|
| ZNF429-gRNA-2 | GAGTTCTGTTGTGCTGTGTCC |
| ZNF429-gRNA-3 | GCTTTCTCTAGACAAGTGATT |
| ZNF257-gRNA-1 | GTAATATGAAGAGACATGAGA |
| ZNF257-gRNA-2 | GCTCATGTCTCTTCATATTAC |
| ZNF257-gRNA-3 | GTTCAGGGACCACTGACAATT |
| ACP1-gRNA-1 | GAAAGCTAAAATTGAACTACT |
| ACP1-gRNA-2 | GCCTGAAAACTGCTTCTGCAA |
| ACP1-gRNA-3 | GTGAGCTGCCTAAGAAATCA |
| LILRA4-gRNA-1 | GCATCAGATACACTCTGTACA |
| LILRA4-gRNA-2 | GAAGAGGGAAACTCAATGTCG |
| LILRA4-gRNA-3 | GAGCCAGGTCCCGTGATCACC |
| ZNF417-gRNA-1 | GCCAGCGAGGATATGAGAGCC |
| ZNF417-gRNA-2 | GCACTGTGACCTTTGAAGATG |
| ZNF417-gRNA-3 | GCGATGGCAGCGGCTGCGCCG |
| IL18R1-gRNA-1 | GCAACAGCACATCATTGTATA |
| IL18R1-gRNA-2 | GAAACAGTTACTATCAAACAC |
| IL18R1-gRNA-3 | GATCATGCAAAGCAATTCTCG |
| NAPSA-gRNA-1 | GTGCTATCAGGTGCAGTATTT |
| NAPSA-gRNA-2 | GCAAACTTGGTCCCATTGGCC |
| NAPSA-gRNA-3 | GGCCCAATATCCCATCAAAA |
| PRAMEF16-gRNA-1 | GCCAGTCCCCATCCAGACTCC |
| PRAMEF16-gRNA-2 | GCTCACCTGGGGCGAAGCTTC |
| PRAMEF16-gRNA-3 | GCATGAGAAAGAGGCAGACAG |
| EPSTI1-gRNA-1 | GAGAATCTGTAAGAATCAAGA |
| EPSTI1-gRNA-2 | GAGCTGAACTCCAAAAAATGA |
| EPSTI1-gRNA-3 | GTGTATTTAGATTGCATCAGC |
| TNFRSF18-gRNA-1 | GTACCCTGGGACTGTACCCCC |
| TNFRSF18-gRNA-2 | GCGAGGAGTGCTGTTCCGAG |
| TNFRSF18-gRNA-3 | GTCGCTCGCCCCGCTCTTCCT |
| IVL-gRNA-1 | GAAGCAATACTTCTCCCTTTG |
| IVL-gRNA-2 | GAAAGCACATGACTGCTGTAA |
| IVL-gRNA-3 | GCTAGCTCTTGATCCAGTTGC |
| PCED1B-gRNA-1 | GAAGTTATGTTTACGTGCCT |
| PCED1B-gRNA-2 | GAGGCCGCTGATAACGGGGTG |
| PCED1B-gRNA-3 | GTTCCGCTCCGACCACCATC |
| HSH2D-gRNA-1 | GAGAGTTTCACTCTGTCACTC |
| HSH2D-gRNA-2 | GTGAAACTCTGTCTCTAAAAT |
| HSH2D-gRNA-3 | GTCCAGTCATTCCCTAGGACT |
| CASC5-gRNA-1 | GTATCAACCAACAGCTTATCT |
| CASC5-gRNA-2 | GATTTATAGAGAAGATTGTG |
| CASC5-gRNA-3 | GAAACATATTGACTTAACATC |
| PRAMEF2-gRNA-1 | GCACGTCAGATAATGAACTCG |
| PRAMEF2-gRNA-2 | GTCTTCATCAGCGATACCAG |
| PRAMEF2-gRNA-3 | GTGCTCTATCTCCCACTCTTC |

| | |
|---|---|
| SPRR2G-gRNA-1 | GCTGCTTGCACTGCTGCTGC |
| SPRR2G-gRNA-2 | GGACAAGGAGGAGGCAGGTA |
| SPRR2G-gRNA-3 | GGATACTTCTGCTGGCAGGG |
| SLAMF6-gRNA-1 | GCACAATTAATCATTCCAAAG |
| SLAMF6-gRNA-2 | GCTCTTTGGAATGATTAATTG |
| SLAMF6-gRNA-3 | GCACGTGGATTTCTGGACTTT |
| ENTPD1-gRNA-1 | GTACCCTTCTGCTGGCTTGTC |
| ENTPD1-gRNA-2 | GAGCTGCAATGAAGGGAACCA |
| ENTPD1-gRNA-3 | GTGCATCAAGTAGAAGAATGC |
| SLC31A2-gRNA-1 | GTGGAGTGTCCACAGTCCTGC |
| SLC31A2-gRNA-2 | GAATTCTTACCAGCAGGACTG |
| SLC31A2-gRNA-3 | GCATCGGCTACTTCATCATGC |
| CXorf27-gRNA-1 | GGCTTGGAGGCCAGCAACAA |
| CXorf27-gRNA-2 | GACCTGTAGCTCATTTCTAGA |
| CXorf27-gRNA-3 | GTGCGCAACACTTCACAAGAT |
| HRNR-gRNA-1 | GGAGTATGATACGTTGAACA |
| HRNR-gRNA-2 | GCGGCCCGAAGCGTGATGGG |
| HRNR-gRNA-3 | GACCATGCTTACTATAGCCAG |
| ZNF141-gRNA-1 | GTTCAGGAACTCTTAACATTC |
| ZNF141-gRNA-2 | GTATCCTCACCCAGGGAGACC |
| ZNF141-gRNA-3 | GCTCTCACCTACCTGGGGGTC |
| GZMB-gRNA-1 | GGCCCACAATATCAAAGAAC |
| GZMB-gRNA-2 | GTCCCCCACGCACAACTCAA |
| GZMB-gRNA-3 | GCATGCCATTGTTTCGTCCAT |
| P2RX5-gRNA-1 | GTTTCAGGGAGCCATTCCTGA |
| P2RX5-gRNA-2 | GACCCCTGCAGGAGTGAAGAC |
| P2RX5-gRNA-3 | GCACCAGGCAAAGATCTCAC |
| ERICH1-gRNA-1 | GGAGGCATCGTGCAATGTTC |
| ERICH1-gRNA-2 | GAACATTGCACGATGCCTCC |
| ERICH1-gRNA-3 | GCGCTGGCAGTGTAGAGCCGT |
| SLC26A3-gRNA-1 | GCAGTAGTGAAGCCACTGATG |
| SLC26A3-gRNA-2 | GCTTACCCACGGATATGTGTC |
| SLC26A3-gRNA-3 | GTTTCCGATTCTGAGTATGA |
| SUSD1-gRNA-1 | GTGTGGCATGTTCATGGCAAG |
| SUSD1-gRNA-2 | GTGGTCGGGTGGAAAGGTTC |
| SUSD1-gRNA-3 | GAACGGGAGGACTCAGTGTGT |
| C1orf127-gRNA-1 | GTCTCAGAAGCATCTTCTTGT |
| C1orf127-gRNA-2 | GATGTGGCTACTTCCTGCATC |
| C1orf127-gRNA-3 | GCTTGTCCGTGTCTGACCTGA |
| C6orf10-gRNA-1 | GTGGGCACGATGTAAGCAAAG |
| C6orf10-gRNA-2 | GCATGCATATTCAACACAAAG |
| C6orf10-gRNA-3 | GTTTGTGTTGAATATGCATGT |
| TSPAN8-gRNA-1 | GTACCCAGAACAAGAAGTTGA |
| TSPAN8-gRNA-2 | GTTACCTTCAACTTCTTGTTC |

| | |
|---|---|
| TSPAN8-gRNA-3 | GTACCCATATTGCTAATGCT |
| C9orf43-gRNA-1 | GACCATATTTCTAACATCTTC |
| C9orf43-gRNA-2 | GTCTCATTCAAATTCAACAC |
| C9orf43-gRNA-3 | GCCTTAGTAAAGGTACATTC |
| ICAM2-gRNA-1 | GTCAGCCGTGCTGTTGAATG |
| ICAM2-gRNA-2 | GCAGGGTCCTGTAACCGAAAG |
| ICAM2-gRNA-3 | GGCTCAACCGCCAGCTTCTT |
| HSD17B6-gRNA-1 | GTCTCCAGCCTGTCAGACGTC |
| HSD17B6-gRNA-2 | GTGCAGCGATGCTCTCCATCT |
| HSD17B6-gRNA-3 | GACTTTGCTTCATTCGCTCTA |
| LILRA3-gRNA-1 | GCAGACTCACCTGCCTGCACG |
| LILRA3-gRNA-2 | GCTCTCTTCCAGGGCTGAGCC |
| LILRA3-gRNA-3 | GTCTCTGAGAGGCCTGCAGTG |
| C3orf17-gRNA-1 | GAGTTTGACTGTGAAACATCT |
| C3orf17-gRNA-2 | GTCACCAGCCCAACCATCACA |
| C3orf17-gRNA-3 | GACCCCATTTGGCCCTCAAAC |
| ZNF679-gRNA-1 | GTGTAGTCATAGAATTCTCTC |
| ZNF679-gRNA-2 | GCTCTCACTTACCTGGGTGTT |
| ZNF679-gRNA-3 | GTCATTTCTCTTTATATTCCA |
| CD86-gRNA-1 | GTCTTTCAGGAACCAACACAA |
| CD86-gRNA-2 | GTTCTTACCAGAGAGCAGGA |
| CD86-gRNA-3 | GAGTAACATTCTCTTTGTGA |
| HMMR-gRNA-1 | GTGAAGCTGCAGGTCACCCAA |
| HMMR-gRNA-2 | GATCTGAAACAGAAAAACTCT |
| HMMR-gRNA-3 | GTTAGAATTCTCAAATTCTTC |
| ATP1B3-gRNA-1 | GATAATTGGATTAAAGCCTGA |
| ATP1B3-gRNA-2 | GAACTGACATGCAACATAAAC |
| ATP1B3-gRNA-3 | GCGTGACCAGATTCCTAGCCC |
| PRR13-gRNA-1 | GCCTCATCCTGTGCCACAGCC |
| PRR13-gRNA-2 | GTTCTTGTCTACTATCACTGC |
| PRR13-gRNA-3 | GTATTCAGTCAGAATCACTGC |
| COL4A2-gRNA-1 | GTGATGGCTGGCCGGGAGCTC |
| COL4A2-gRNA-2 | GATTGGACTGCCTGGTGGCAA |
| COL4A2-gRNA-3 | GACCCGGCTTCCTATGAATCC |
| COX5A-gRNA-1 | GCACTGACCTTAACAACCTCT |
| COX5A-gRNA-2 | GATGAGGAGTTTGATGCTCGC |
| COX5A-gRNA-3 | GACCACCCGGGCCGACCCTCG |
| CBFB-gRNA-1 | GCCTTTGAAGAGGCTCGGAGA |
| CBFB-gRNA-2 | GGAGGATGCATTAGCACAAC |
| CBFB-gRNA-3 | GTTTCTAAGTCGACATACTCT |
| CD300C-gRNA-1 | GTCTGAGCCACCCCATGACCG |
| CD300C-gRNA-2 | GTCTCCGATGTGACAAGATTG |
| CD300C-gRNA-3 | GGGTTCGGGGCTGTCCTTTC |
| PECR-gRNA-1 | GATCCTGCAGTTTACAGCTCC |

| | |
|---|---|
| PECR-gRNA-2 | GTTCCCATACAATGCAACATC |
| PECR-gRNA-3 | GATCTTACCAAAAGTATCTA |
| C17orf66-gRNA-1 | GCCAGAACTCAGAAAAGCCA |
| C17orf66-gRNA-2 | GTCTGCCCTTGTCCTGCTACC |
| C17orf66-gRNA-3 | GATAGAACTGCTCATCCTCTC |
| FLG-gRNA-1 | GTCATTACGAGTTTGTCTGC |
| FLG-gRNA-2 | GTCATCTGCAGTCAGCGATCG |
| FLG-gRNA-3 | GATTGAGTAAAAAAGAGCTGA |
| ANKRD36-gRNA-1 | GTACCTGCCAAGATAATCAA |
| ANKRD36-gRNA-2 | GTCACAGGACCGCCCTACATT |
| ANKRD36-gRNA-3 | GCTACTATATACCTTGATCAG |
| FPR1-gRNA-1 | GATTCGTGTGACTACAGTACC |
| FPR1-gRNA-2 | GCCTGGATATCATCACTTATC |
| FPR1-gRNA-3 | GCCCATAACTGACAGCAACGA |
| PODXL-gRNA-1 | GCACAGAGGGTGTTTCCTGTG |
| PODXL-gRNA-2 | GTCGTGGGCTGCACTGTCTCC |
| PODXL-gRNA-3 | GGTGTTCTCAATGCCGTTGC |
| ITGB3BP-gRNA-1 | GAAATTGTCAGAAGAAATCA |
| ITGB3BP-gRNA-2 | GAAAATTCTTACTTAGTTCTT |
| PSG2-gRNA-1 | GACAGCTGCTATGTTGGATTA |
| PSG2-gRNA-2 | GAGCTGAAACCTATGAGTCAT |
| PSG2-gRNA-3 | GTCGCGAAGCAAGACAAGTAG |
| IFIT3-gRNA-1 | GCTGCCTCGTTGTTACCATCT |
| IFIT3-gRNA-2 | GTCCTGATAACCAATACGTCA |
| IFIT3-gRNA-3 | GGCAATTGCGATGTACCATC |
| HERC5-gRNA-1 | GAAAATAATTCAGATCACATG |
| HERC5-gRNA-2 | GCATTCTCTTGCACTCTCAAA |
| HERC5-gRNA-3 | GACTATAGCATGAAACATCTA |
| CLEC4A-gRNA-1 | GCTAAATGACTTCCAATTCTT |
| CLEC4A-gRNA-2 | GAGAATGAGATTGCCAATAGC |
| CLEC4A-gRNA-3 | GTGCCAATGTCGCTGACCTTC |
| CXCL9-gRNA-1 | GTCCTTCACATCTGCTGAATC |
| CXCL9-gRNA-2 | GCCCAGATTCAGCAGATGTGA |
| CXCL9-gRNA-3 | GCCAGCAAGATGATGCCCAAG |
| CENPO-gRNA-1 | GTGGAAGTATTATCCACTTCA |
| CENPO-gRNA-2 | GCTCATTGATCTCCACTGTT |
| CENPO-gRNA-3 | GCAGCTCTTCAGACTGTTTA |
| CD300A-gRNA-1 | GCTGTCCCAGAACCCCAAGC |
| CD300A-gRNA-2 | GAAGCAGCAACAATGCCAGC |
| CD300A-gRNA-3 | GGGCCTCCCTGCTAGCCTGG |
| MEFV-gRNA-1 | GCCTGAGCGCCAATCAGCTC |
| MEFV-gRNA-2 | GATGCGACCTAGAAGCCTTG |
| MEFV-gRNA-3 | GCTGGCAGCTCCGCCCCCGTA |
| GCSAML-gRNA-1 | GAATCAAAAGAAGCCCAAGAA |

| | |
|---|---|
| GCSAML-gRNA-2 | GACCCAGATGAGGAAAGAAAA |
| GCSAML-gRNA-3 | GTCTACTTACCTGATTAGAAG |
| SERPING1-gRNA-1 | GAATAGCAAGAAGTACCCTG |
| SERPING1-gRNA-2 | GCTTTGGTCGTGAAGCCCTTC |
| SERPING1-gRNA-3 | GCAACAACAGTGACGCCAACT |
| NDUFB2-gRNA-1 | GCACCTTCAGTCTTCATCATC |
| NDUFB2-gRNA-2 | GCGTATCCTGATCCTTCCCAG |
| NDUFB2-gRNA-3 | GTTGTCTTGCAGTGCCGGTGG |
| CAMP-gRNA-1 | GCACACACTAGGACTCTGTCC |
| CAMP-gRNA-2 | GACGACACAGCAGTCACCAG |
| CAMP-gRNA-3 | GGGGACAGTGACCCTCAACC |
| CD33-gRNA-1 | GTGCTACTGCTGCCCCTGCTG |
| CD33-gRNA-2 | GCCACTCACCTGCCCACAGC |
| CD33-gRNA-3 | GTTCCTACCTGAGCCATCTCC |
| SLAMF1-gRNA-1 | GAACCATTACCAGACAACAG |
| SLAMF1-gRNA-2 | GGATGCAGGACAGACCCCTC |
| SLAMF1-gRNA-3 | GTATGCCCAAGTCCAGAAACC |
| CD209-gRNA-1 | GCGTCGTAATCAAAAGTGCTG |
| CD209-gRNA-2 | GTCTTGTATCCTCGAGTCTGT |
| CD209-gRNA-3 | GCATCACCGCCTGCAAAGAAG |
| NPIP-gRNA-1 | GTCCAGGAATTAAAATTGTCC |
| NPIP-gRNA-2 | GCACATCTTACTGTTCTTTGG |
| NPIP-gRNA-3 | GACTATCTTCCCGTCTCAAAA |
| TSLP-gRNA-1 | GACCTGATTACATATATGAG |
| TSLP-gRNA-2 | GCCTTAGCTATCTGGTGCCC |
| TSLP-gRNA-3 | GCACCCGATTGCTACAAGAGA |
| IRF7-gRNA-1 | GTCCGGATGCAGGCGAGGCCG |
| IRF7-gRNA-2 | GGACCCGGCCGACCCGCACA |
| IRF7-gRNA-3 | GATGCACTCACCTTGCACCG |
| MyD88-gRNA-1 | GCTCCCCTAGGTGCCGCCGGA |
| MyD88-gRNA-2 | GTGGGGATCAGTCGCTTCTGA |
| MyD88-gRNA-3 | GTCTACAGCGGCCACCTGTAA |
| TLR2-gRNA-1 | GGAACTGCGAGATACTGATT |
| TLR2-gRNA-2 | GTGTGACATTCCGACACCGAG |
| TLR2-gRNA-3 | GGTTCAGGATGTCCGCCTCT |
| TLR7-gRNA-1 | GCCTGCGGTATCTCTAGTAGC |
| TLR7-gRNA-2 | GTGCCTTCCAGTTGCGATATC |
| TLR7-gRNA-3 | GCGCAAGCTCACCCATACTTC |
| TMEM173-gRNA-1 | GGATGTTCAGTGCCTGCGAG |
| TMEM173-gRNA-2 | GCTGATTGTAAGTTCGAATC |
| TMEM173-gRNA-3 | GGTGCCTGATAACCTGAGTA |
| NONO-gRNA-1 | GCACGAGCCTCCTTGATGTTG |
| NONO-gRNA-2 | GCTGCATACTTGTGAAATTGC |
| NONO-gRNA-3 | GAGCTGCACGCCATGAGCACC |

| | |
|---|---|
| XPO1-gRNA-1 | GCTTGATTTCAGCCAAAAAC |
| XPO1-gRNA-2 | GAAGTATTTGATTTCTCTAG |
| XPO1-gRNA-3 | GTTAAATGCTTAGATTTGACT |
| DDX58-gRNA-1 | GTTTGAAATCCCAACTTTCAA |
| DDX58-gRNA-2 | GTCTGAATGTTTAATTAATC |
| DDX58-gRNA-3 | GATGCCTTCTCAGATCAGACA |
| IRF3-gRNA-1 | GGCACCAACAGCCGCTTCAG |
| IRF3-gRNA-2 | GGGAGTGGGATTGTCCAAGC |
| IRF3-gRNA-3 | GATCTACGAGTTTGTGAACTC |
| PQBP1-gRNA-1 | GCAAGCGGGTCTGCAGCGCAA |
| PQBP1-gRNA-2 | GACAGCGGGCTCCCTTACTAC |
| PQBP1-gRNA-3 | GACAAGGTGTTCGACCCTTCC |
| DDX3X-gRNA-1 | GAACTCTTCAGATAATCAGAG |
| DDX3X-gRNA-2 | GTCTAGCTTCTTCAGTGATCG |
| DDX3X-gRNA-3 | GACCTACCTTTAGTAGCTTCT |
| TLR8-gRNA-1 | GAAGTCAAGCAACTCGAGACG |
| TLR8-gRNA-2 | GAGAGGTTTAGGTATCGAAGT |
| TLR8-gRNA-3 | GAATCTGTTGAGCGTCGTTTC |
| TLR9-gRNA-1 | GCCGCAAGACGCTGTTTGTGC |
| TLR9-gRNA-2 | GCTCCGAAGCTCCGCTGATG |
| TLR9-gRNA-3 | GCCGGACTGCCACACTTCACC |
| IFITM1-gRNA-1 | GAAGCCAGAAGATGCACAAGG |
| IFITM1-gRNA-2 | GTTCATCCTGTTACTGGTATT |
| IFITM1-gRNA-3 | GACAGCACCAGTTCAAGAAG |
| IFITM2-gRNA-1 | GCCGCTGTTGACAGGAGAGA |
| IFITM2-gRNA-2 | GGAGCAGGAAGTGGCTATGC |
| IFITM2-gRNA-3 | GCTCCTTGAGCATCTCGTAGT |
| IFITM3-gRNA-1 | GCATACGCACCTTCACGGAGT |
| IFITM3-gRNA-2 | GTTCTTCTCTCCTGTCAACAG |
| IFITM3-gRNA-3 | GGAGCACGAGGTGGCTGTGC |
| EZR-gRNA-1 | GCAATCCAGCCAAATACAAC |
| EZR-gRNA-2 | GACGCAGGTGGTAAAGACTAT |
| EZR-gRNA-3 | GCAATGTCCGAGTTACCACCA |
| SERINC5-gRNA-1 | GTCCAAAACAGACTCTATACA |
| SERINC5-gRNA-2 | GAGCTTTCTTCATTCCAGATC |
| SERINC5-gRNA-3 | GCCGTGTATAGAGTCTGTTT |
| DLG1-gRNA-1 | GTACGCTTGTATGTAAAAAGA |
| DLG1-gRNA-2 | GTGCTCCCCCTGTGATAATTT |
| DLG1-gRNA-3 | GAAGCTCATTAAAGGTCCTAA |
| SERINC3-gRNA-1 | GCCTGAGCTGAAATAGCCCCC |
| SERINC3-gRNA-2 | GTAGGAACACCAGCCTCGCTC |
| SERINC3-gRNA-3 | GAACAAGTAAAGATCTCCGAG |
| MARCH8-gRNA-1 | GTTCTATCACGCCATCCAGCC |
| MARCH8-gRNA-2 | GCGTGATAGAAGTGCGAGAGA |

| | |
|---|---|
| MARCH8-gRNA-3 | GAAGCCTCCACTTCGTGCACC |
| SELPLG-gRNA-1 | GACAACTCGACTGACGGCCA |
| SELPLG-gRNA-2 | GCCTGCTGCAAGGCGTTCTAC |
| SELPLG-gRNA-3 | GTGGTCTAGAAGGAGATAAGA |
| MAP3K5-gRNA-1 | GCCGAGTTAGTATCACAGTAG |
| MAP3K5-gRNA-2 | GTGGGAGGCCAAATCAAAGGT |
| MAP3K5-gRNA-3 | GCTATGATTCTATTGTGAAGC |
| SUMO1-gRNA-1 | GAGAATCATACTGTCAAAGAC |
| SUMO1-gRNA-2 | GTCCCCCAAGTCCTCAGTTGA |
| SUMO1-gRNA-3 | GGAGGCAAAACCTTCAACTG |
| CTR9-gRNA-1 | GCTGATAACTTCATCTCCCTC |
| CTR9-gRNA-2 | GAATAATATTCCAGCCCTTCT |
| CTR9-gRNA-3 | GAGCCTGCTTCTGCCTACTTG |
| TRIM37-gRNA-1 | GCAAACATGAAGAAAATGAAA |
| TRIM37-gRNA-2 | GCTCCCCAAAGTGCACACTGA |
| TRIM37-gRNA-3 | GTCTGCCATCAGTGTGCACTT |
| RAD18-gRNA-1 | GTTACAGTTATGTGAACACTG |
| RAD18-gRNA-2 | GAGACAATAGATGATTTGCTG |
| RAD18-gRNA-3 | GACAGTGGATTGTCCTGTTTG |
| SAMHD1-gRNA-1 | GCGTTCACTTATCTGCAGCTC |
| SAMHD1-gRNA-2 | GAATCCACGTTGATACAATGA |
| SAMHD1-gRNA-3 | GTCTTCGATACATCAAACAGC |
| APOBEC3D-gRNA-1 | GATCTGGAAGCGCCTGTTAGC |
| APOBEC3D-gRNA-2 | GTGAACATGAATCCACAGATC |
| APOBEC3D-gRNA-3 | GATTGGCGGTGGGTGCTCCTC |
| APOBEC3H-gRNA-1 | GATTCCAGGGGTACGTGCGC |
| APOBEC3H-gRNA-2 | GCGACCCTGCGCACGTACCCC |
| APOBEC3H-gRNA-3 | GAAGTCACCTGTTACCTCACG |
| SUN2-gRNA-1 | GCGTCCCTGAAGACGTTCCTC |
| SUN2-gRNA-2 | GGACAGGCGCTTCATGTTGC |
| SUN2-gRNA-3 | GCCCACTCACGTCGTCCCGCA |
| SUN1-gRNA-1 | GCTCCTGTATTGGACGAGTCT |
| SUN1-gRNA-2 | GTCGTGGCCAGGCGCAAACTA |
| SUN1-gRNA-3 | GTCGAGTCACAGCATCCTGC |
| TRIM28-gRNA-1 | GTTGCACATAACCAGATCGCC |
| TRIM28-gRNA-2 | GCCTTGGTGTACTTCACCCGC |
| TRIM28-gRNA-3 | GATTGAGCTGGCAGTCTCGGC |
| CDKN1A-gRNA-1 | GAGTCGAAGTTCCATCGCTCA |
| CDKN1A-gRNA-2 | GTGTACCCTTGTGCCTCGCTC |
| CDKN1A-gRNA-3 | GATTTCTACCACTCCAAACGC |
| CEBPB-gRNA-1 | GAGGTAAGCGCGCAGCGCGTA |
| CEBPB-gRNA-2 | GTGAAGTCGATGGCGCGCTCG |
| CEBPB-gRNA-3 | GCGATGTTGTTGCGCTCGCGC |
| DDX5-gRNA-1 | GTAGCACTTACCAGGGAAAT |

| | |
|---|---|
| DDX5-gRNA-2 | GACCCACTGCTATTCAAGCTC |
| DDX5-gRNA-3 | GTCCCTGAGCTTGAATAGCAG |
| HSPA1A-gRNA-1 | GCCTGGGGGCTTCGGGGCTC |
| HSPA1A-gRNA-2 | GCCCCTAATCTACCTCCTCAA |
| HSPA1A-gRNA-3 | GCCCTGAGCCCCGAAGCCCCC |
| HSPA1B-gRNA-1 | GCCCCTAATCCACCTCCTCAA |
| HSPA1B-gRNA-2 | GCACCGGCATGGCCAAAGCCG |
| HSPA1B-gRNA-3 | GCCTGGCGGCTTCGGGGCTC |
| HSPA6-gRNA-1 | GTCATGAAGCCGAGCAGTACA |
| HSPA6-gRNA-2 | GATCCTCAGCCTTGTACTGCT |
| HSPA6-gRNA-3 | GCTGCGAGTCATTGAAATAGG |
| RBPJ-gRNA-1 | GTTTCCAGGAAATTTGGTGAG |
| RBPJ-gRNA-2 | GGGGAAAGATGGGGGGCTGC |
| RBPJ-gRNA-3 | GTCTCAACCGTGTGCATTTAT |
| CIITA-gRNA-1 | GTCCTACCTGTCAGAGCCCCA |
| CIITA-gRNA-2 | GCCCCTAGAAGGTGGCTACC |
| CIITA-gRNA-3 | GCATCGCTGTTAAGAAGCTCC |
| GADD45B-gRNA-1 | GAGGCAGAGGACCACGCTGTC |
| GADD45B-gRNA-2 | GCATGACGCTGGAAGAGCTCG |
| GADD45B-gRNA-3 | GACTCGTACACCCCCACTGTG |
| NFKBIA-gRNA-1 | GCCAAAAGCTCCACGATGCCC |
| NFKBIA-gRNA-2 | GTGCTGCTGTATCCGGGTGCT |
| NFKBIA-gRNA-3 | GTGTCAACAGAGTTACCTACC |
| POU2F2-gRNA-1 | GGAGTCCAGACCTTGCTTCT |
| POU2F2-gRNA-2 | GCAGGTGCTTACCTTTGTACT |
| POU2F2-gRNA-3 | GTGCGGTAGCAGGAACTGAGC |
| ARHGEF1-gRNA-1 | GTGGCCCTGCAGTTTGAGCC |
| ARHGEF1-gRNA-2 | GCGGCGCCCAGCCCACCTCA |
| ARHGEF1-gRNA-3 | GCTCAGCCCCGATGATGCTGA |
| TRIM22-gRNA-1 | GTTATCACTAGGAGTGAAAGC |
| TRIM22-gRNA-2 | GATGTGATTGACGTCATGAAA |
| TRIM22-gRNA-3 | GTCAGACAAGAGAGAACCGCC |
| SLC40A1-gRNA-1 | GTGTGATTGCAGTAGCAGTAC |
| SLC40A1-gRNA-2 | GAACATTCTGTACCACCAGCG |
| SLC40A1-gRNA-3 | GCTACTGCAATCACAATCCAA |
| CNP-gRNA-1 | GTCTCCCGGCAGTCGTCCCTG |
| CNP-gRNA-2 | GCCCGTAGTACCCCGTGAAGA |
| CNP-gRNA-3 | GTTTGTGACACCCAAGACGAC |
| FOXO1-gRNA-1 | GGTGGCGCAAACGAGTAGCA |
| FOXO1-gRNA-2 | GGCTGCCAACCCCGACGCCG |
| FOXO1-gRNA-3 | GTAGGTTGACATGACCGAATT |
| HSPA9-gRNA-1 | GTGCGAGTCATTGAAATAAGC |
| HSPA9-gRNA-2 | GTTCTGATTTGTCTAGACCAT |
| HSPA9-gRNA-3 | GCTCTAGGCCACTAAAGATGC |

| | |
|---|---|
| SMARCA1-gRNA-1 | GTGAGTAAGATGCAACGAGAA |
| SMARCA1-gRNA-2 | GAAGAGTCTGCCACCTAAAA |
| SMARCA1-gRNA-3 | GCTTCTTGCTCTGTGCGCCTA |
| SUPT6H-gRNA-1 | GTTGGGTGTCAAAGTCAAAAG |
| SUPT6H-gRNA-2 | GCAAAAAAATGTCAGATGACG |
| SUPT6H-gRNA-3 | GCACCCAAATTCTCCTCAATG |
| TRAF6-gRNA-1 | GAGTTGACAATGAAATACTGC |
| TRAF6-gRNA-2 | GCCATGACCAGAACTGTCCTT |
| TRAF6-gRNA-3 | GTTTGGAAGGGACGCTGGCAT |
| UBP1-gRNA-1 | GATGGAGCTTCACAGACCTC |
| UBP1-gRNA-2 | GCTCCATCTCCCTGATGATT |
| UBP1-gRNA-3 | GTCTGTCCTTAGTGTTCTCCG |
| DENND4A-gRNA-1 | GATACTTGTGTTATAAAAAAT |
| DENND4A-gRNA-2 | GAACACTGTATCTTACAAAGC |
| DENND4A-gRNA-3 | GCTGACTACTTTGTTGTAGC |
| DNAJC5-gRNA-1 | GAGTCATTGTACCACGTCCT |
| DNAJC5-gRNA-2 | GTGTAAGCCCAAGGCGCCTGA |
| DNAJC5-gRNA-3 | GAACTCCGTCTCCTCGCCTTC |
| CTNNB1-gRNA-1 | GCTCCTTCTTGATGTAATAAA |
| CTNNB1-gRNA-2 | GAGTGAAGGCGAACTGCATTC |
| CTNNB1-gRNA-3 | GAAAATGGCAGTGCGTTTAGC |
| SUPT4H1-gRNA-1 | GGTCTTTATAGCTGTGTCTC |
| SUPT4H1-gRNA-2 | GTGCAGTCATATACCATCTCT |
| SUPT4H1-gRNA-3 | GTGATGCATATCTACAAATGA |
| HEXIM1-gRNA-1 | GTTCCATGAAGTCGTCATCGC |
| HEXIM1-gRNA-2 | GAAGTGCCTCTCGCGCATGG |
| HEXIM1-gRNA-3 | GCGAGCGCCGCTTTCCAAGTT |
| EHMT2-gRNA-1 | GTGAGTTCAGCTTCCTCCTTT |
| EHMT2-gRNA-2 | GTTCTTCCAGATCTGGCCAAA |
| EHMT2-gRNA-3 | GAGGGGGCTGACACCCCTGT |
| TCF4-gRNA-1 | GTTTGACAAATTACTCAAAGA |
| TCF4-gRNA-2 | GAATCTAAGAACATTATCTCT |
| TCF4-gRNA-3 | GAAGAGATAATGTTCTTAGAT |
| BRD4-gRNA-1 | GTAAGATCATTAAAACGCCTA |
| BRD4-gRNA-2 | GACAGGAGGAGGATTCGGCTG |
| BRD4-gRNA-3 | GTCGATGCTTGAGTTGTGTT |
| GADD45A-gRNA-1 | GCTGGAGAGCAGAAGACCGAA |
| GADD45A-gRNA-2 | GATCCATGTAGCGACTTTCC |
| GADD45A-gRNA-3 | GCAGTGATCGTGCGCTGACTC |
| MYC-gRNA-1 | GTCGCTTACCAGAGTCGCTGC |
| MYC-gRNA-2 | GCGCCGTCGTTGTCTCCCCGA |
| MYC-gRNA-3 | GACAACGTCTTGGAGCGCCAG |
| PRDX2-gRNA-1 | GAGGGGCCTCTTTATCATCGA |
| PRDX2-gRNA-2 | GTAATCCTCAGACAAGCGTC |

| | |
|---|---|
| PRDX2-gRNA-3 | GCGTGACCAGACGCTTGTCTG |
| GADD45G-gRNA-1 | GCACTTACACGTTCAAGACTT |
| GADD45G-gRNA-2 | GCAGCGGCTGGCGGCTATCG |
| GADD45G-gRNA-3 | GCACCCAGTCGTTAACGCTG |
| PRMT6-gRNA-1 | GCCTCTACCGCGTACACGCGC |
| PRMT6-gRNA-2 | GCCCATCCACTCGCTCACGA |
| PRMT6-gRNA-3 | GTCCTCCACGCGCGAACCAAG |
| UBASH3B-gRNA-1 | GCACAGAAGCTTTCCGACTTT |
| UBASH3B-gRNA-2 | GCAGAACATTGACGTCAAGCT |
| UBASH3B-gRNA-3 | GGAGGTAGAGGACGTACTCC |
| BIRC2-gRNA-1 | GGCTTGAGGTGTTGGGAATC |
| BIRC2-gRNA-2 | GATATCCTCATCTTCTTGAAC |
| BIRC2-gRNA-3 | GCATGGGTAGAACATGCCAAG |
| RHOA-gRNA-1 | GAGCAAGCATGTCTTTCCAC |
| RHOA-gRNA-2 | GAGCCGGTGAAACCTGAAGA |
| RHOA-gRNA-3 | GTATCGAGGTGGATGGAAAGC |
| CHD1-gRNA-1 | GACCTCACTTGATGAGTCAT |
| CHD1-gRNA-2 | GTTCCGATGACTCATCAAGTG |
| CHD1-gRNA-3 | GAGCCAGTCGGATGATGATTC |
| CHD3-gRNA-1 | GTCAAAGATAAGGATGACATT |
| CHD3-gRNA-2 | GTTCTTCACACCCAATGCTGA |
| CHD3-gRNA-3 | GTCTAAGATGATGACCATCCT |
| DKC1-gRNA-1 | GTGAATCCAAAGTTGCTAAGT |
| DKC1-gRNA-2 | GCGTGTCCAACTTAGCAACTT |
| DKC1-gRNA-3 | GTAGGCCCTAGAAACTCTGAC |
| E2F1-gRNA-1 | GCTGCTGAGCCACTCGGCTGA |
| E2F1-gRNA-2 | GCGCGCATGGGCCGCCCGCAG |
| E2F1-gRNA-3 | GCGACGACACCGTCAGCCGAG |
| GRN-gRNA-1 | GCTGGAACGCGGTGCCCAGA |
| GRN-gRNA-2 | GCAGCTGCTGCCGTCCCCTTC |
| GRN-gRNA-3 | GCGATCCTGCTTCCAAAGATC |
| HDAC1-gRNA-1 | GCACCATGCAAAGAAGTCCG |
| HDAC1-gRNA-2 | GTTACGTCAATGATATCGTCT |
| HDAC1-gRNA-3 | GCTGGATACGGAGATCCCTAA |
| HDAC2-gRNA-1 | GCCAGAACACTCCAGAATATA |
| HDAC2-gRNA-2 | GAAATATGGGGAATACTTTCC |
| HDAC2-gRNA-3 | GTTCTGGTTTGTCATGTTTGA |
| HMGB1-gRNA-1 | GAACATCCTGGCCTGTCCAT |
| HMGB1-gRNA-2 | GAGTATCGCCCAAAAATCAA |
| HMGB1-gRNA-3 | GAGAAGTTGACTGAAGCATC |
| HSPA4-gRNA-1 | GTTAATGAATGAAACCACTGC |
| HSPA4-gRNA-2 | GCATGGCAGTCACTTGCTCAG |
| HSPA4-gRNA-3 | GACTGCACAATATCATATGCA |
| HSPA8-gRNA-1 | GATGCCTGGGGGATTTCCTGG |

| | |
|---|---|
| HSPA8-gRNA-2 | GAGATCGATTCTCTCTATGA |
| HSPA8-gRNA-3 | GTTTAAGCGCAAGCATAAGA |
| HSPB1-gRNA-1 | GTGTATTTCCGCGTGAAGCAC |
| HSPB1-gRNA-2 | GTTACTTGGCGGCAGTCTCAT |
| HSPB1-gRNA-3 | GAACTTGGGTGGGGTCCACAC |
| MCM2-gRNA-1 | GCAACAAGTGCAATTTCGTCC |
| MCM2-gRNA-2 | GAGTCCTCATAGTTCACCACC |
| MCM2-gRNA-3 | GAACCAGCTGATCCGCACCAG |
| MECP2-gRNA-1 | GCCGCCGCCGCCGCGCCGAG |
| MECP2-gRNA-2 | GAACAGGATTCCATGGTAGCT |
| MECP2-gRNA-3 | GATGATGGAGCGCCGCTGTT |
| PRDX1-gRNA-1 | GCCCAGCGCTCACTTCTGCT |
| PRDX1-gRNA-2 | GGCTTGATGGTATCACTGCC |
| PRDX1-gRNA-3 | GACTGTAAATGACCTCCCTGT |
| POU2F1-gRNA-1 | GAGAAACCAGTAAACCATCTA |
| POU2F1-gRNA-2 | GAGGAGCAATCTCAACAGCCC |
| POU2F1-gRNA-3 | GTTGAGATTGCTCCTCCTAC |
| SMARCA4-gRNA-1 | GGCGTGCCCCCCGGGATGCC |
| SMARCA4-gRNA-2 | GCGGCACCTCCAAATTACAGC |
| SMARCA4-gRNA-3 | GTTGTCCTGAGGGTACCCTCC |
| SP3-gRNA-1 | GCGAAAAGCCCGTGAAACAAG |
| SP3-gRNA-2 | GGGACTTACCGCTCCGGCGG |
| SP3-gRNA-3 | GCCGCAACTTGACCAAGTGTG |
| SUPT5H-gRNA-1 | GAGGCGAAGCTTCGGAAGAGA |
| SUPT5H-gRNA-2 | GCAAAGCGGTTGTCCTTCTTC |
| SUPT5H-gRNA-3 | GCACGATGACACCCACAGTC |
| TFCP2-gRNA-1 | GGAGAACTTCCAGAAATTAA |
| TFCP2-gRNA-2 | GCACTGAGCATCAGCAGCTAG |
| TFCP2-gRNA-3 | GTGTACTGAAGCCTTCTGTCA |
| NELFA-gRNA-1 | GAGACCGCCCAGCAGTTGAAG |
| NELFA-gRNA-2 | GGGAGTGAGGGGTCCCGCGA |
| NELFA-gRNA-3 | GTATGGTTGGCGCTGGCCGAG |
| ARID1A-gRNA-1 | GAATACTCACAGGCAAGCTGG |
| ARID1A-gRNA-2 | GTGAGCGAGACTGAGCAACAC |
| ARID1A-gRNA-3 | GTTCAATAGATGACCTCCCCA |
| SETDB1-gRNA-1 | GTAACTTACCTTCTGGTCTTT |
| SETDB1-gRNA-2 | GCAGAACTCCAAAAGACCAGA |
| SETDB1-gRNA-3 | GCCTAACCTTTATGCAGATCC |
| DNAJB6-gRNA-1 | GTATGACAAATATGGCAAAGA |
| DNAJB6-gRNA-2 | GCAGAGACATGCCTCACCCG |
| DNAJB6-gRNA-3 | GTGGATTACTATGAAGTTCT |
| GNA13-gRNA-1 | GTTGAAACAATCGTCAATAAC |
| GNA13-gRNA-2 | GAGATGATGTCGTTTGATACC |
| GNA13-gRNA-3 | GGAACTCCTCGCGCGCGCGC |

| | |
|---|---|
| CBX3-gRNA-1 | GAGAGCCTGAAGAATTTGTCG |
| CBX3-gRNA-2 | GAAACAGGAAAGATTCAGATG |
| CBX3-gRNA-3 | GAGAAACGCTTCAATCAATTC |
| TRIM32-gRNA-1 | GCCAGTTTGTAGTAACCGATG |
| TRIM32-gRNA-2 | GAGAGAAGTTAACTCGTCTG |
| TRIM32-gRNA-3 | GCTATAGTGTCCTTATTCGAG |
| DICER1-gRNA-1 | GCAATCCACCACAATCTCACA |
| DICER1-gRNA-2 | GAGCCCACTTCTGTCAGTAAA |
| DICER1-gRNA-3 | GTCCATCATGTCCTCGCATTT |
| NELFB-gRNA-1 | GACTCCTGAAGCAGTACATCC |
| NELFB-gRNA-2 | GAACGTGAAGCTGTACGACA |
| NELFB-gRNA-3 | GAGCCTTCCCCTCCGAAGCGA |
| DROSHA-gRNA-1 | GTTTGCGCTGCATTTGCAGAG |
| DROSHA-gRNA-2 | GACCAAAGTTCATCATGAAGT |
| DROSHA-gRNA-3 | GACTCTGCAAATGCAGCGCAA |
| EIF3L-gRNA-1 | GCACCTACCATTGCCAACCTG |
| EIF3L-gRNA-2 | GAGCGACTATGATATGCACAC |
| EIF3L-gRNA-3 | GATTCTTCAAGAATACACCT |
| NELFCD-gRNA-1 | GACAGTTTCCTGAACCTGCAC |
| NELFCD-gRNA-2 | GCTGCAAATCTAACTTCATCA |
| NELFCD-gRNA-3 | GTTCAGGTGTTGAGCCAGTGC |
| BANP-gRNA-1 | GCAACCTCCAGATCCATCACG |
| BANP-gRNA-2 | GACCGTCCTGCCCCACGTGA |
| BANP-gRNA-3 | GATGGTGGCCGGCTCCCCTCT |
| ACTL6A-gRNA-1 | GTTACTCACAGTGTGTTATCC |
| ACTL6A-gRNA-2 | GTCTCAGGAAGCTGTTCGTGA |
| ACTL6A-gRNA-3 | GTACCGAAGCTTGAAAATCC |
| CYLD-gRNA-1 | GTAGAACCTTTGCTAAAAATA |
| CYLD-gRNA-2 | GAGAATATTCAAGAATTCCTC |
| CYLD-gRNA-3 | GACTTAGACTCAACCTTATTC |
| HIF1A-gRNA-1 | GTTATGGTTCTCACAGATGA |
| HIF1A-gRNA-2 | GTTTAAATGAGCTCCCAATGT |
| HIF1A-gRNA-3 | GTACTCATCCATGTGACCATG |
| DNAJA1-gRNA-1 | GCACCATTATATAAGTCTTCT |
| DNAJA1-gRNA-2 | GATATGACAAAGGAGGAGAAC |
| DNAJA1-gRNA-3 | GTTAGAAGTTCATATTGACAA |
| HSPA2-gRNA-1 | GCCGCCTCACGGCGCGCTTGT |
| HSPA2-gRNA-2 | GACCCCGACGCACGAATAGG |
| HSPA2-gRNA-3 | GATCTCGATGCTCGCCTGCG |
| HSPA5-gRNA-1 | GTAAAATGAAAGAAACCGCTG |
| HSPA5-gRNA-2 | GCGTTGGGCATCATTAAAAT |
| HSPA5-gRNA-3 | GCGCCAAGCAACCAAAGACGC |
| DNAJB1-gRNA-1 | GCATCTCTCTTAAAGATATTG |
| DNAJB1-gRNA-2 | GGCTTCACCAACGTGAACTT |

| | |
|---|---|
| DNAJB1-gRNA-3 | GCCCAAAACACCCGAGAAACG |
| PRKAA1-gRNA-1 | GTTTCAGAAATCTGATGTACC |
| PRKAA1-gRNA-2 | GAGTAAAAACAGGCTCCACGA |
| PRKAA1-gRNA-3 | GTTGGCAAACATGAATTGAC |
| HSPA13-gRNA-1 | GTACTGACAATGATGTATATG |
| HSPA13-gRNA-2 | GAGTCGAGAGCCAACATATTC |
| HSPA13-gRNA-3 | GGCTGACGTCTTCCACGTCT |
| YY1-gRNA-1 | GTGAGGGCAAGCTATTGTTCT |
| YY1-gRNA-2 | GAAAACTAAAACGACACCAAC |
| YY1-gRNA-3 | GCCTCTCAGATCCCAAACAAC |
| ZNF10-gRNA-1 | GATTCACCAAGAGACCCATCC |
| ZNF10-gRNA-2 | GTAGACACTGGTGACCTTCA |
| ZNF10-gRNA-3 | GATGTATTTGTGGACTTCACC |
| EIF3F-gRNA-1 | GTTTAGCCCCAACAGAGTGAT |
| EIF3F-gRNA-2 | GCATATTGCAACACTGTACTC |
| EIF3F-gRNA-3 | GCAGCCCAGGATGAGCTCATT |
| HDAC3-gRNA-1 | GAGCAACCCAGCTGAACAACA |
| HDAC3-gRNA-2 | GCAGACCACCAGCCCAGTTAA |
| HDAC3-gRNA-3 | GCCCTATGAAGCCCCATCGCC |
| MTA1-gRNA-1 | GCTCTGTGGGCACCTTCGCA |
| MTA1-gRNA-2 | GATAAAGGAGAGATTCGAGT |
| MTA1-gRNA-3 | GATCACCGACTTGTTAAAAGA |
| PRDX4-gRNA-1 | GCTGAAGTTAACTGATTATCG |
| PRDX4-gRNA-2 | GCCAACTGAAATTATCGCTTT |
| PRDX4-gRNA-3 | GGTGTATACCTAGAGGACTC |
| FAM208A-gRNA-1 | GATCAACCATCAGAAATGCAT |
| FAM208A-gRNA-2 | GACTGCCAAGAATTGTTATTT |
| FAM208A-gRNA-3 | GTTCAAACCATATGTGAAAA |
| TARDBP-gRNA-1 | GATTCTGCATGCCCCAGATGC |
| TARDBP-gRNA-2 | GTCAAGAAAGATCTTAAGAC |
| TARDBP-gRNA-3 | GCCCATGGAAAACAACCGAAC |
| HspBP1-gRNA-1 | GGGCCCCCTCTCGCTCTTGC |
| HspBP1-gRNA-2 | GTTGGTCTCCAGAGGCGTCAG |
| HspBP1-gRNA-3 | GGGCTCCAAGCACGTGCTCG |
| LEF1-gRNA-1 | GTGGATACTTACACGTGCATT |
| LEF1-gRNA-2 | GAGGACCGGGAATCATATGA |
| LEF1-gRNA-3 | GTTACTTACTGTTCTCGGGA |
| HSPA14-gRNA-1 | GAATGAAAGTAAAGCAGATCC |
| HSPA14-gRNA-2 | GTGATTCCATACCTTCTGCCC |
| HSPA14-gRNA-3 | GCGATGTATTTCTGAGCTTG |
| PPHLN1-gRNA-1 | GCCACCACTGCTAGACAGACC |
| PPHLN1-gRNA-2 | GCTTCCCATTATGCGAGAGAG |
| PPHLN1-gRNA-3 | GCTTACACTGGGATGACTTCG |
| CPSF3-gRNA-1 | GAGTGGAGCTGGGCAAGAAGT |

| | |
|---|---|
| CPSF3-gRNA-2 | GCTACAGAAGACAAGTTTCAA |
| CPSF3-gRNA-3 | GCATCCATTCCTTCTAGGCC |
| HEXIM2-gRNA-1 | GCCGCCGACGGTGTTTCTTCC |
| HEXIM2-gRNA-2 | GGTTCTCCTGACGAAGCCGC |
| HEXIM2-gRNA-3 | GACTTACCTTGGCCTCCTCC |
| COMMD1-gRNA-1 | GCTGACAGCGTCTTCAGAATT |
| COMMD1-gRNA-2 | GTACATCATCTTCAGTTAGGC |
| COMMD1-gRNA-3 | GCCTAATTCCAGCTCTATAA |
| MST1R-gRNA-1 | GAGCCAGGACACTCCTTCTGC |
| MST1R-gRNA-2 | GAGCCATACCTCAGTAAGCTT |
| MST1R-gRNA-3 | GATAGCAGTGCAACCCCTCTT |
| NELFE-gRNA-1 | GTCTCATACAGAGATTTCCTC |
| NELFE-gRNA-2 | GAGTCATCCAGACGTCCCCAG |
| NELFE-gRNA-3 | GCTCAGCCTTGATGGCACTGA |
| RNF7-gRNA-1 | GATTGTTCTGTTTCACCCACA |
| RNF7-gRNA-2 | GTCAAGCTGAAAACAAACAAG |
| RNF7-gRNA-3 | GACAACTGCTGCATGTCCCTG |
| SIRT1-gRNA-1 | GCAACTTTGTATTTACAAATC |
| SIRT1-gRNA-2 | GACCATTCTTCAAGTTTGCAA |
| SIRT1-gRNA-3 | GACCTTTGCAAACTTGAAGAA |
| TRIM11-gRNA-1 | GACTGGTAGAGACACTGCGG |
| TRIM11-gRNA-2 | GACATCAAGGACGCCCTGCGC |
| TRIM11-gRNA-3 | GACCTGCCTGCCACAAGACGC |
| HSPA12A-gRNA-1 | GTCTGTCCCCCTCCCATATTG |
| HSPA12A-gRNA-2 | GACTTCTTACCACAATATGGG |
| HSPA12A-gRNA-3 | GCGGAATGCATCCATGTGATG |
| BCL11A-gRNA-1 | GTCACAGATAAACTTCTGCAC |
| BCL11A-gRNA-2 | GATCTACTTAGAAAGCGAACA |
| BCL11A-gRNA-3 | GTGTTTATCAACGTCATCTAG |
| CAV1-gRNA-1 | GTTTAGGGTCGCGGTTGACC |
| CAV1-gRNA-2 | GTGGGGGCAAATACGTAGACT |
| CAV1-gRNA-3 | GACAGACGGTGTGGACGTAGA |
| HMOX1-gRNA-1 | GCAGAGCCCCTCACGGGCACC |
| HMOX1-gRNA-2 | GAAGGGCCAGGTGACCCGAGA |
| HMOX1-gRNA-3 | GCGAAGCCCTGGTGCCCGTG |
| LIF-gRNA-1 | GCTTGGCCTTCTCCGTGCCGT |
| LIF-gRNA-2 | GCAGCCCATAATGAAGGTCT |
| LIF-gRNA-3 | GAGCCAACTGGCACAGCTCAA |
| MMP8-gRNA-1 | GCTATACCCCACAGCTGTCAG |
| MMP8-gRNA-2 | GTGCAACACTCCAGAGTTCAA |
| MMP8-gRNA-3 | GATTCCATTGGGTCCATCAAA |
| PRKAA2-gRNA-1 | GTTATCTAATATGATGTCAGA |
| PRKAA2-gRNA-2 | GTACCTGCCTGAGATGACTTC |
| PRKAA2-gRNA-3 | GAATGCCAAGATAGCCGATTT |

| | |
|---|---|
| SUV39H1-gRNA-1 | GCGTGTGTTGCAAGTCTTCT |
| SUV39H1-gRNA-2 | GGATCTTCTTGTAATCGCAC |
| SUV39H1-gRNA-3 | GTCGCAAGAACAGCTTCGTCA |
| TFAP4-gRNA-1 | GACAGCTCAAGCGCTTCATCC |
| TFAP4-gRNA-2 | GAGCCGAGTACATCTTCTCCC |
| TFAP4-gRNA-3 | GCACGATCACCGTGGGGTGGT |
| ZNF175-gRNA-1 | GAATTTATCCCAGAAGCCTC |
| ZNF175-gRNA-2 | GTATCACATTCCCAACCCAG |
| ZNF175-gRNA-3 | GCTCACCCACTGCGAAGAGA |
| MTA2-gRNA-1 | GCATTACCAGCCACCCACATA |
| MTA2-gRNA-2 | GACGGATTGAGGAGCTCAACA |
| MTA2-gRNA-3 | GTGTCCTGTACCGTATGTGGG |
| IKBKE-gRNA-1 | GATGGGCAGGAGCTAATGTTT |
| IKBKE-gRNA-2 | GATGCAGACGGACTCTGGAAG |
| IKBKE-gRNA-3 | GTTCCAGAGTCCGTCTGCATG |
| FOXP3-gRNA-1 | GTGCCCCCCAGCTCTCAACGG |
| FOXP3-gRNA-2 | GCCTGGACACCCATTCCAGGC |
| FOXP3-gRNA-3 | GCCACTTACAGGCACTCCTCC |
| UBASH3A-gRNA-1 | GCTCCGCCGTCTTCCTCCCCG |
| UBASH3A-gRNA-2 | GCCTAGACGACCCCATCCCCC |
| UBASH3A-gRNA-3 | GCTTCTGCAGGCTGAAAGCGT |
| ZNF350-gRNA-1 | GCACCTGAACAGGCTCCACTG |
| ZNF350-gRNA-2 | GACTCCTGGGCGCTGCTCAGA |
| ZNF350-gRNA-3 | GCAACTGTGGACAATTGAAGA |
| BCL11B-gRNA-1 | GTGAGTCCCGTCACCCGAGAC |
| BCL11B-gRNA-2 | GGCAATGTCCCGCCGCAAAC |
| BCL11B-gRNA-3 | GCCAGACCCTCGTCTTCTTCG |
| HSPA12B-gRNA-1 | GGGCAATGCCGCAGCTTTCC |
| HSPA12B-gRNA-2 | GCACTGGGGACCGCTCCGGGC |
| HSPA12B-gRNA-3 | GCCGCGATTACTACCATGACC |
| DUSP1-gRNA-1 | GCCATGGCTGTGCTCGACCG |
| DUSP1-gRNA-2 | GTGCGTATCACGCTTCCCGCA |
| DUSP1-gRNA-3 | GTAATCGAGTCAAGCTGGACG |
| ISG15-gRNA-1 | GCACGCCTTCCAGCAGCGTC |
| ISG15-gRNA-2 | GCGACGAACCTCTGAGCATCC |
| ISG15-gRNA-3 | GTGTGCAGGACGACCTGTTC |
| AXIN1-gRNA-1 | GCCGGTACTTACGGGAATGTG |
| AXIN1-gRNA-2 | GGGTGTCTGCATCGCTGGAC |
| AXIN1-gRNA-3 | GCCCCAAGGTCTTCGGGGACG |
| CDK13-gRNA-1 | GTCCTGATTTGCTTGAAATAT |
| CDK13-gRNA-2 | GATTGTATAGCTCAGAAGAA |
| CDK13-gRNA-3 | GCAGATAAAACTTGCAGACTT |
| EIF3E-gRNA-1 | GTATTCTGATGATATTCCTCA |
| EIF3E-gRNA-2 | GCACTAATTTAGAATCAATCT |

| | |
|---|---|
| EIF3E-gRNA-3 | GTGCATTTGCCTTGTAGTTTC |
| SFPQ-gRNA-1 | GTCATCCTCCGTGATATCAGC |
| SFPQ-gRNA-2 | GTGAAATTGCCAAAGCCGAAC |
| SFPQ-gRNA-3 | GCGTACTCAAACGTGCCATGC |
| DGCR8-gRNA-1 | GATCATGACATTCCATAACTC |
| DGCR8-gRNA-2 | GCCTACAGAGCCGCTGCCCGA |
| DGCR8-gRNA-3 | GGATGACTTTGACAACGATG |
| BST2-gRNA-1 | GTACATTAAACCATAAGCTTC |
| BST2-gRNA-2 | GCTCTTCTTAGATGGCCCTAA |
| BST2-gRNA-3 | GCGATTCTCACGCTTAAGACC |
| ZC3H12A-gRNA-1 | GTCTTACGCAGGAAGTTGTCC |
| ZC3H12A-gRNA-2 | GCGGCCCGACGTGCCCATCAC |
| ZC3H12A-gRNA-3 | GAATCGGCACTTGATCCCAT |
| SLFN11-gRNA-1 | GCTGAACACCCTGGGACCATT |
| SLFN11-gRNA-2 | GAGGTATTTCCTGAAGCCGAA |
| SLFN11-gRNA-3 | GCAGCCTGACAACCGAGAAA |
| ZAP70-gRNA-1 | GCGGGCTCATCTACTGCCTGA |
| ZAP70-gRNA-2 | GAGCAGTGCCAGCAACGCCTC |
| ZAP70-gRNA-3 | GGAGAAGCTCATTGCTACGA |
| MOV10-gRNA-1 | GCCGGATCACCGGAAACCCTG |
| MOV10-gRNA-2 | GCAGCGCTAAGGGCTATGACC |
| MOV10-gRNA-3 | GCCGACTCCAGGTCATAGTGC |
| TNFRSF10D-gRNA-1 | GATTTACAAACTGTACATAGC |
| TNFRSF10D-gRNA-2 | GCAACAGAGGCGCAGCCTCA |
| TNFRSF10D-gRNA-3 | GTCTCATAGATCAGAATATAC |
| ADAR-gRNA-1 | GCATGGGTGTCGTCGTCAGCT |
| ADAR-gRNA-2 | GTGCATACACTCAAGCAGTG |
| ADAR-gRNA-3 | GAGCCATGACAATTCTGCTAG |
| DDX6-gRNA-1 | GCTTGTATATTGTCCTTCTTC |
| DDX6-gRNA-2 | GTGAAACCCACTGGTGGCCC |
| DDX6-gRNA-3 | GTAACATACCTGAATAGGAGA |
| CD164-gRNA-1 | GACCTGATGTAGTAACTGTCT |
| CD164-gRNA-2 | GAACAGTTAGTGATTGTCAAG |
| CD164-gRNA-3 | GTCCAAGACAGTTACTACATC |
| LSM1-gRNA-1 | GTCATCAAGAGTATCTGCTCG |
| LSM1-gRNA-2 | GCTGACCACAAAAATCCCTCG |
| LSM1-gRNA-3 | GCGAGATGGAAGGACACTTAT |
| TNRC6A-gRNA-1 | GTTTGAAAACTGTTTAACAAA |
| TNRC6A-gRNA-2 | GTTTACAAAGGGAATGATGAA |
| PRKRA-gRNA-1 | GAGAATATACTACAATTTGC |
| PRKRA-gRNA-2 | GCTTACCTGTAATGAACCAAT |
| PRKRA-gRNA-3 | GAAGAACCAGCTTAATCCTAT |
| ROCK2-gRNA-1 | GAAAATTGTGAAAAAAATCAG |
| ROCK2-gRNA-2 | GTATGATGTTGTAAAAGTTAT |

| | |
|---|---|
| ROCK2-gRNA-3 | GAAATCAACAGGTTCGTCACA |
| XRN1-gRNA-1 | GTTTCACCACTTCGCTGAGAC |
| XRN1-gRNA-2 | GTATCCCTGTCTCAGCGAAG |
| XRN1-gRNA-3 | GTAATGCGAAACAACACCTCC |
| AGO2-gRNA-1 | GGGGCCGGCTCCCGAGTACA |
| AGO2-gRNA-2 | GACCCTTTGGGAATCTGTCAG |
| AGO2-gRNA-3 | GTCACAAACAAACTCGATTAC |
| RNASEL-gRNA-1 | GCAGATCACCCACAGTGTTC |
| RNASEL-gRNA-2 | GTGCTCTTATCAAAATCTGCC |
| RNASEL-gRNA-3 | GGACTGTCTGAGTGACCTGC |
| TNFRSF10A-gRNA-1 | GACACACTCGATGTCACTCCA |
| TNFRSF10A-gRNA-2 | GACAGCATGTCAGTGCAAACC |
| TNFRSF10A-gRNA-3 | GCAGGCAATGGACATAATATA |
| TSPO-gRNA-1 | GTTTCAGGTACGGCTCCTACC |
| TSPO-gRNA-2 | GCGGGACAACCATGGCTGGCG |
| TSPO-gRNA-3 | GCTTCTTTGGTGCCCGACAAA |
| IFI30-gRNA-1 | GCAAACCCCAGGCCTGCGTGT |
| IFI30-gRNA-2 | GTCAAGTTCATCCAACACGC |
| IFI30-gRNA-3 | GCACCACACGAGTATGTGCCC |
| LGALS3BP-gRNA-1 | GCTGGTGTGGTCTGCACCAA |
| LGALS3BP-gRNA-2 | GCCAGATCTTTGACAGCCAG |
| LGALS3BP-gRNA-3 | GTCCATCAGCGTGAATGTGC |
| HSP90AB1-gRNA-1 | GCATACTCACATAAAGGGTGA |
| HSP90AB1-gRNA-2 | GAGATACTCACTGAGATACTC |
| HSP90AB1-gRNA-3 | GACAAGAGCCTCACTAATGAC |
| DHX30-gRNA-1 | GTGCAGCCTGCCAGCTGTTCA |
| DHX30-gRNA-2 | GCCTTCTACCTCCACGCTCT |
| DHX30-gRNA-3 | GCTAGTCTACGTGCACACAAA |
| RNF115-gRNA-1 | GAGGCCCTCCCCCAGCTGACA |
| RNF115-gRNA-2 | GTCCAACAGTGACAGTAACTC |
| RNF115-gRNA-3 | GAATAAAGCCTGATTCACATC |
| CAV2-gRNA-1 | GCAGATCCACACTTTGTCAA |
| CAV2-gRNA-2 | GAATGGCCAGGAACACCGTC |
| CAV2-gRNA-3 | GGCGGACGTACAGCTCTTCA |
| TRIM21-gRNA-1 | GTGTGCAGCAAAAAAACTTCC |
| TRIM21-gRNA-2 | GTCAGGAGGTGATAATTGTCC |
| TRIM21-gRNA-3 | GTCAGTTCCCCTAATGCCACC |
| CCDC8-gRNA-1 | GTCGACTATTTGCTCATCCCC |
| CCDC8-gRNA-2 | GCACTGGCAGGTTGACGCGG |
| CCDC8-gRNA-3 | GCTCCTTTCGGCGCCGCAGGA |
| RSAD2-gRNA-1 | GACAACCTTCTGGTTAGATTC |
| RSAD2-gRNA-2 | GAAAGACTCCTACCTTATTC |
| RSAD2-gRNA-3 | GAGCGTGAGCATCGTGAGCAA |
| CD37-gRNA-1 | GACCAGCTTCGTGTCCTTTG |

| | |
|---|---|
| CD37-gRNA-2 | GATTCCCAGGGTGATCTGTG |
| CD37-gRNA-3 | GTTTGCCACACAGATCACCC |
| CD53-gRNA-1 | GTGTACTCACAAATGACTGGA |
| CD53-gRNA-2 | GGCACGAGTGATTGGACCAG |
| CD53-gRNA-3 | GTGCCCAGCTGAATGAGTATG |
| HAVCR2-gRNA-1 | GTATAGCAGAGACACAGACAC |
| HAVCR2-gRNA-2 | GCTTACTGTTAGATTTATATC |
| HAVCR2-gRNA-3 | GAGGTCACCCCTGCACCGACT |
| CD63-gRNA-1 | GCATCTGCCTGCATCCTGTCC |
| CD63-gRNA-2 | GCTTTCTGTCTCTTATCATGT |
| CD63-gRNA-3 | GTCTAAACACATAGCCAGCAA |
| CD81-gRNA-1 | GCGGCCACCTCACAGGCAAAC |
| CD81-gRNA-2 | GCATGATGTTCGTTGGCTTCC |
| CD81-gRNA-3 | GTGACAGCGCCCACAGCGATG |
| CHMP5-gRNA-1 | GTGACAAGAAGATTTCTCGAT |
| CHMP5-gRNA-2 | GTCGAGAAATCTTCTTGTCAA |
| CHMP5-gRNA-3 | GATGAAGATGATTTAGAAGC |
| CD151-gRNA-1 | GCCACAGCCCCAATGACCCTC |
| CD151-gRNA-2 | GCGTCGGAACCTGCTGCGCC |
| CD151-gRNA-3 | GCTGGATGAAGGTCTCCAACT |
| ABCA1-gRNA-1 | GTCCTGATCCTGATCTCTGTT |
| ABCA1-gRNA-2 | GTTTGCGCATGTCCTTCATGC |
| ABCA1-gRNA-3 | GAGCTCAGCCGAACAGAGATC |
| CD9-gRNA-1 | GCACCAGCATCATGAGGGCGC |
| CD9-gRNA-2 | GCTGAAAGCCATCCACTATG |
| CD9-gRNA-3 | GATGGCCGCAGCTATTTCAA |
| HGS-gRNA-1 | GCCTGGCGCTGTCACAGTCAG |
| HGS-gRNA-2 | GCACTCTCTCGGCAGCAAACA |
| HGS-gRNA-3 | GCCTCTGACTGTGACAGCGCC |
| UBE2L6-gRNA-1 | GCATTGGCATCATCGCTGGAC |
| UBE2L6-gRNA-2 | GCAAGATCTACCACCCCAACG |
| UBE2L6-gRNA-3 | GAAGAGTTCACCCTCCGATT |
| CD82-gRNA-1 | GATCCTGGGCGCAGTGATCC |
| CD82-gRNA-2 | GGATGCCTGGGACTACGTGC |
| CD82-gRNA-3 | GGCCCTCTTCTACTTCAACA |
| TSG101-gRNA-1 | GTCACCATATGAATCCAAAAC |
| TSG101-gRNA-2 | GACAATCCCTGTGCCTTATAG |
| TSG101-gRNA-3 | GTGACAGTTTCACGTACAGTT |
| CC2D1B-gRNA-1 | GTCTCCCTAGACGGAGCTGC |
| CC2D1B-gRNA-2 | GAGTTGGCGGCAGACTGTATG |
| CC2D1B-gRNA-3 | GAAGACCTCGGAAGGGCCCTC |
| TIMD4-gRNA-1 | GCCAAAGAACCTCTCATTCTC |
| TIMD4-gRNA-2 | GATCATCAGCCAGAGAATGAG |
| TIMD4-gRNA-3 | GTTTACATCGTTGAACCAGCC |

| | |
|---|---|
| TSPAN7-gRNA-1 | GTAGTTTCATGGAGACTAACA |
| TSPAN7-gRNA-2 | GATCATCGCTGGAGTGGCGTT |
| TSPAN7-gRNA-3 | GAGGGTTTGTGTTTCGTCATG |
| UBA7-gRNA-1 | GACCTGCCATGCAGAGGATTC |
| UBA7-gRNA-2 | GACCCATCAGAACCAAGTTCT |
| UBA7-gRNA-3 | GCTTTCTGGCGGCTGACACC |
| CLEC4M-gRNA-1 | GCCACGTGCCTTCCTGATTT |
| CLEC4M-gRNA-2 | GAAGAAGATCCAACAACCAG |
| CLEC4M-gRNA-3 | GCGTCGTAATCAAAACTGCTG |
| CC2D1A-gRNA-1 | GGAGGCTGATGATGACCTGC |
| CC2D1A-gRNA-2 | GCCAAACTGTACCTGGGCCAC |
| CC2D1A-gRNA-3 | GCCTGCAGGCGGAGCTAAATG |
| KCNK3-gRNA-1 | GCGCCGCGGTCTTCGACGCGC |
| KCNK3-gRNA-2 | GCTCCACGTCCGACACGTGCG |
| KCNK3-gRNA-3 | GCGGAGCCCGAGCTGATCGAG |
| CFLAR-gRNA-1 | GCTCACCCTGAAGTTATTTGA |
| CFLAR-gRNA-2 | GTTTCTCCAACTCAACCACA |
| CFLAR-gRNA-3 | GTACCAGACTGCTTGTACTTC |
| XIAP-gRNA-1 | GCATTTAGCATGTTGTTCCCA |
| XIAP-gRNA-2 | GCAAATATCTGTTAGAACAGA |
| XIAP-gRNA-3 | GCCAATTCTTCTAGTTAGTGA |
| BCL2L1-gRNA-1 | GTGCCCGGGAGGTGATCCCCA |
| BCL2L1-gRNA-2 | GCAGGCGACGAGTTTGAACTG |
| BCL2L1-gRNA-3 | GAGACCCCCAGTGCCATCAA |
| KAT5-gRNA-1 | GAAGCGAAAATCGAATTGTTT |
| KAT5-gRNA-2 | GGCCGAGATCCTGAGCGTGA |
| KAT5-gRNA-3 | GACTCACCTGGGGAAAAACCG |
| RPL12-gRNA-1 | GCTGGGCCTGTCTGTTCTGAA |
| RPL12-gRNA-2 | GTTCTAGTCTCCAAAAAAAGT |
| RPL12-gRNA-3 | GTCTTCTGCAGTTAAACACAG |
| BTG2-gRNA-1 | GCCCCTCCAAGAACTACGTGA |
| BTG2-gRNA-2 | GCTCCTCGTACAAGACGCAGA |
| BTG2-gRNA-3 | GGTCTTCAGCGGGGCGCTCC |
| AKAP13-gRNA-1 | GCAATTCCTAGCAACCAGTGC |
| AKAP13-gRNA-2 | GACATATAAGGGAGCTTGCTG |
| AKAP13-gRNA-3 | GTCCGTCACTGTACAAGTACT |
| STAB1-gRNA-1 | GCCTGCTTGGATGGCATGGAC |
| STAB1-gRNA-2 | GTTCGGGCCTGACTGCCAATC |
| STAB1-gRNA-3 | GCCTGTCCATGCCATCCAAGC |
| TAGLN2-gRNA-1 | GCATTAATGCACTGTACCCCG |
| TAGLN2-gRNA-2 | GCTGGCAGTAGCCCGAGATGA |
| TAGLN2-gRNA-3 | GCAGCGGACGCTGATGAATC |
| MID1IP1-gRNA-1 | GGGCACGTCGCGCAGCAAGC |
| MID1IP1-gRNA-2 | GCCAACATCGTTGTCTAACC |

| | |
|---|---|
| MID1IP1-gRNA-3 | GAACGCCATGAATCGCTTCAT |
| RBM10-gRNA-1 | GCACGGGAGGTTCGGCTGATG |
| RBM10-gRNA-2 | GCCGCCGCCTACGCTGAGTCT |
| RBM10-gRNA-3 | GCACGCCGTGCGACTGCAGC |
| PSEN2-gRNA-1 | GCACACAGGGGGCGTGAAGCT |
| PSEN2-gRNA-2 | GTGTAGAAGATGAAGTCCCCG |
| PSEN2-gRNA-3 | GATGACCCACGCGGACCACTC |
| DHX9-gRNA-1 | GGAAGAACAAGAAGTGCAAG |
| DHX9-gRNA-2 | GCCTCGGTCCCAGGTGGGCCC |
| DHX9-gRNA-3 | GCCCCACTGCTCTAATTTCAT |
| DHX15-gRNA-1 | GTGGTGTAATAATTCTTGATG |
| DHX15-gRNA-2 | GAGTTGCCTGTACCCAACCC |
| DHX15-gRNA-3 | GGAGTATAGAAGATCTCAAC |
| FPGS-gRNA-1 | GCTGAACATCATCCACGTCAC |
| FPGS-gRNA-2 | GTCCCCTTCGTCCCAGTGACG |
| FPGS-gRNA-3 | GTAGAGGCGCCAGAAGTACT |
| MDM2-gRNA-1 | GTACCAAGTTCCTGTAGATCA |
| MDM2-gRNA-2 | GAGACACTTATACTATGAAAG |
| MDM2-gRNA-3 | GCTTGGTAGTAGTCAATCAGC |
| POLR2A-gRNA-1 | GTGACTCACACGGGGCAAGCC |
| POLR2A-gRNA-2 | GACATGTCTACGACCTTTGCA |
| POLR2A-gRNA-3 | GCAAAAACATATGCGAGGGT |
| RAB6A-gRNA-1 | GTCCGCTGAGGAAATTCAAGC |
| RAB6A-gRNA-2 | GTTTATCAAAAACTATGTACT |
| RAB6A-gRNA-3 | GTGTAACTGGGAATGAGGCTA |
| RAD21-gRNA-1 | GAGGTGTGGTTGACCTGCCTG |
| RAD21-gRNA-2 | GATCGTGAGATAATGAGAGA |
| RAD21-gRNA-3 | GACATACTCTAAGTCAGGCAG |
| SRSF6-gRNA-1 | GAACCTATGCGGATGCCCACA |
| SRSF6-gRNA-2 | GTACAGCAGTCGGAGAACATC |
| SRSF6-gRNA-3 | GATAGGTCTCGATCTAGAAGA |
| UBE2B-gRNA-1 | GTATGACTTACCATCTTCAAA |
| UBE2B-gRNA-2 | GTAGATATCCTTCAGAATCGA |
| UBE2B-gRNA-3 | GTCTTTGCAGTGTATGCTGA |
| SLU7-gRNA-1 | GCAAAAAGAAAGACTGCTTTG |
| SLU7-gRNA-2 | GTACAAGAGGGGTGTAAAAG |
| SLU7-gRNA-3 | GCAAAAGCAGTTCAGCTCATC |
| NCKAP1-gRNA-1 | GTTACAATATCAAAGAAGACT |
| NCKAP1-gRNA-2 | GTATTCAGCAATTCACAAACA |
| NCKAP1-gRNA-3 | GTGCTGTCTCGAATTGAAGAA |
| NCOA6-gRNA-1 | GTTCGGTTCTGAGCCAAAGCC |
| NCOA6-gRNA-2 | GTTTGAGACTAAATTCTGCGA |
| NCOA6-gRNA-3 | GTCGCAGAATTTAGTCTCAA |
| R3HDM1-gRNA-1 | GACCACAATGTTGATCAGAG |

| | |
|---|---|
| R3HDM1-gRNA-2 | GTACATCTTGGGAAAATATT |
| R3HDM1-gRNA-3 | GCTTTGACAAAGATGATAACC |
| RNF216-gRNA-1 | GAGATTTCCAGATGTAGCAAA |
| RNF216-gRNA-2 | GCATATCTGACTCCTCAGATG |
| RNF216-gRNA-3 | GATAAACCCATTTGCTACATC |
| XAB2-gRNA-1 | GCAGACGGTGGACCCCTTCA |
| XAB2-gRNA-2 | GTCCGTTGTCCTCATAAAACT |
| XAB2-gRNA-3 | GGCGCAGCAAGACGCTGTTG |
| RBM25-gRNA-1 | GCTTAGTACCCACTGTGTCTA |
| RBM25-gRNA-2 | GCTTGTACTCACAGAATCCGA |
| RBM25-gRNA-3 | GTCCTGTACCAATGAGCATTA |
| ELAVL1-gRNA-1 | GAGAGCGATCAACACGCTGAA |
| ELAVL1-gRNA-2 | GTGTACCACTCGCCAGCGCGA |
| ELAVL1-gRNA-3 | GTCCTCGTGGATCAGACTAC |
| SERPINB6-gRNA-1 | GTGTTACCGTTCTCAAGTCAG |
| SERPINB6-gRNA-2 | GCGAGACCACTGACTTGAGAA |
| SERPINB6-gRNA-3 | GCGTGCCAGTCTTGTTCACTT |
| PPIA-gRNA-1 | GAGAGAAAGGATTTGGTTATA |
| PPIA-gRNA-2 | GAAGGTTGGATGGCAAGCATG |
| PPIA-gRNA-3 | GCGTACCTGACACATAAACCC |
| PSMB2-gRNA-1 | GCAACTTTATAAGATGCGAAA |
| PSMB2-gRNA-2 | GTTTCAGCTATCTCACGTGAG |
| PSMB2-gRNA-3 | GTGTGAAGTTAGCTGCTGCCG |
| ST3GAL3-gRNA-1 | GACAGTTTAGAGTCCAGATTC |
| ST3GAL3-gRNA-2 | GACCACCGTGCAGAGAGTCCT |
| ST3GAL3-gRNA-3 | GTGAGCCTAGTGTTTGTCCAG |
| SUMO2-gRNA-1 | GCAACGATCATATTAATTTGA |
| SUMO2-gRNA-2 | GCATGGCCGACGAAAAGCCCA |
| SUMO2-gRNA-3 | GTGAAACAGACACACCTGCAC |
| SNRPC-gRNA-1 | GTGAGAAAGACACACTGCAG |
| SNRPC-gRNA-2 | GTGAAAGACTATTATCAGAAA |
| TOP1-gRNA-1 | GAACACAAAGATCGAGAACAC |
| TOP1-gRNA-2 | GTGAAAAGAAACACAAAGAGA |
| TOP1-gRNA-3 | GTGAGCTTCCATCTTTGTGTT |
| Gemin2-gRNA-1 | GTGAACAAACATAGAAGTCAC |
| Gemin2-gRNA-2 | GTTGACCCAAAGAAGTTGAAA |
| Gemin2-gRNA-3 | GATTTGAGCTACCACAACATC |
| CHST1-gRNA-1 | GGTCGCCCCGCGTGTTGTTC |
| CHST1-gRNA-2 | GGGCTACAAGATCGCCGCCT |
| CHST1-gRNA-3 | GCGGCGATCTTGTAGCCCAGC |
| RNF10-gRNA-1 | GCAACCTCCTGTGGTCTCTA |
| RNF10-gRNA-2 | GTCTAGGTAGCAGAGGCTCAA |
| RNF10-gRNA-3 | GATCCTGATACATTAGTTAAC |
| RBM5-gRNA-1 | GTGCTCAGTGGTCATCCACCC |

| | |
|---|---|
| RBM5-gRNA-2 | GATCTACCTACCTGAGCATAT |
| RBM5-gRNA-3 | GTACCCCTAGCGATCATTCTT |
| PRPF8-gRNA-1 | GCCAGGGAATCTCATTGACGA |
| PRPF8-gRNA-2 | GCTGCGTGGGGCATGTACTTT |
| PRPF8-gRNA-3 | GTCATAGAACCAGTCCAACAC |
| SUB1-gRNA-1 | GCCTGAAGGTGAAATGAAACC |
| SUB1-gRNA-2 | GTAATTGATATTAGAGAATAT |
| SUB1-gRNA-3 | GCAGAGATGATAACATGTTTC |
| SPEN-gRNA-1 | GTAACTTCAGTGCAGATACA |
| SPEN-gRNA-2 | GAAAGCCTTGACTGCATCAAA |
| SPEN-gRNA-3 | GTCTTGGAAATAATCGCCTCA |
| RBM17-gRNA-1 | GCCTCAGGATCCTGTTCCCAG |
| RBM17-gRNA-2 | GATGAAGATTATGAGCGAGAG |
| RBM17-gRNA-3 | GACTCCACCGCATGTAGCAGC |
| AKAP1-gRNA-1 | GCCTCGATCTCCCAGATGATG |
| AKAP1-gRNA-2 | GCATCACCGTGGAGGTCATTG |
| AKAP1-gRNA-3 | GTGTGTGCTGCTGCACGAAC |
| MED7-gRNA-1 | GGATGATAAGATCATCACAT |
| MED7-gRNA-2 | GTCGATGCCCTGACTTTCCAA |
| MED7-gRNA-3 | GATGATAGCAACAATTGTAC |
| BMP2K-gRNA-1 | GTAAGACTCCAATAATTCAC |
| BMP2K-gRNA-2 | GATACCTGTGAAGCTGTTGCA |
| BMP2K-gRNA-3 | GCAAAAATATTGTGGGCTATT |
| RANBP17-gRNA-1 | GTTGAAAGACATGTTGCTGCA |
| RANBP17-gRNA-2 | GTCCTTTACCTGTTGAGCAG |
| RANBP17-gRNA-3 | GTTTGAAGAGTTTGGCTGAAT |
| ERCC5-gRNA-1 | GATTATTTGAAGCAATGCCAG |
| ERCC5-gRNA-2 | GAAATAATGTTCTTCTGCGCT |
| ERCC5-gRNA-3 | GAGATGAAGGGGGCTTTCTGA |
| AP1G2-gRNA-1 | GCACCCTCTTGTCTGTGAATC |
| AP1G2-gRNA-2 | GCCTGGGGGCCATGCTTCTAT |
| AP1G2-gRNA-3 | GCAGAATCCTCAATCCTAACA |
| NUMBL-gRNA-1 | GCCCATCGGCTGACACCCAC |
| NUMBL-gRNA-2 | GTCACGCTGGCCGCCGCGCTG |
| NUMBL-gRNA-3 | GCCCTGGATGTCCCGCAGCG |
| PPP1R14D-gRNA-1 | GTCTCTCAGGCTTATTTCTG |
| PPP1R14D-gRNA-2 | GCATCCCTTACCTCTGTGGGG |
| PPP1R14D-gRNA-3 | GTCAGTCAACTCAAGAAACTC |
| NEIL3-gRNA-1 | GCTTGAGTATAAATATAAAAA |
| NEIL3-gRNA-2 | GTTGACTCATCAGTAGAACTC |
| NEIL3-gRNA-3 | GTTATCTTACCGTAAAGCTTT |
| ZNF587-gRNA-1 | GCACTGTGACCTTTGAAGATG |
| ZNF587-gRNA-2 | GCCAGCGAGGATATGAGAGCC |
| ZNF587-gRNA-3 | GATTACCTGAGTTGGGCGCCT |

| | |
|---|---|
| KPNA4-gRNA-1 | GTGGAGAAGCCTCAGGAAAAG |
| KPNA4-gRNA-2 | GATGTCTGTGAGCAAGCAGTG |
| IPO7-gRNA-1 | GAGCAAATTCATTATACTCCT |
| IPO7-gRNA-2 | GATTTGCTGAAGTATTTCTGA |
| IPO7-gRNA-3 | GCACTTCTTGCATTTCCACCA |
| HMGA1-gRNA-1 | GTGCCAACACCTAAGAGACCT |
| HMGA1-gRNA-2 | GCTCGAAGTCCAGCCAGCCCT |
| HMGA1-gRNA-3 | GCTCACCGGAGGCTGCTTGCG |
| RANBP2-gRNA-1 | GTGTAAAAATGATGTTACTGA |
| RANBP2-gRNA-2 | GACCTTTAGTTTATAAATTGC |
| RANBP2-gRNA-3 | GACTTACCCTGTAACATTCAA |
| NUP210-gRNA-1 | GCACCAGCATCATCTTCGCAG |
| NUP210-gRNA-2 | GGTGGCCAGCATCGAGCCGC |
| NUP210-gRNA-3 | GCACCACCCGCGAGCTCTACC |
| NUP160-gRNA-1 | GCAGTCTTGCTGTTCATTGTG |
| NUP160-gRNA-2 | GTACATGCATGCACCAAAACG |
| NUP160-gRNA-3 | GTAATGCAACGATTGCTTAC |
| NUP54-gRNA-1 | GTCATTCAAAGGACTCTTTGG |
| NUP54-gRNA-2 | GTCGAAGCCAACAACAACAGT |
| NUP54-gRNA-3 | GTGCCCTCTACATTTACAGTA |
| NUP133-gRNA-1 | GAACTATGAAGCTATTAAAGA |
| NUP133-gRNA-2 | GATTTGGACTTGAAGCAAAAC |
| NUP133-gRNA-3 | GCAGCTGAAGTTCTTTGCAAA |
| NUP85-gRNA-1 | GAAACCTCCTTCAACAAAAA |
| NUP85-gRNA-2 | GTCACATACCAGCATCTCCCC |
| NUP85-gRNA-3 | GTACATGCTCTGATGACTGAT |
| KPNA2-gRNA-1 | GAGCGGAGAAGTAGCATCATC |
| KPNA2-gRNA-2 | GACCTGGCAGCTTGAGTAGCT |
| KPNA2-gRNA-3 | GACATCAGTGAACAAGCTGTC |
| NUP155-gRNA-1 | GTAGTGAGACTATTCTTGCTG |
| NUP155-gRNA-2 | GCTCACTAAGTCCATCAAAAT |
| NUP155-gRNA-3 | GAGCAAACCAGGTCCTTGCAA |
| CPSF6-gRNA-1 | GAAAAAACTTACCCCTTTGAC |
| CPSF6-gRNA-2 | GGAGAGTATCCATGTAATCT |
| CPSF6-gRNA-3 | GCTTCAGATCCAACACCAACA |
| TNPO3-gRNA-1 | GTGTTTGCACACATCCCTTCC |
| TNPO3-gRNA-2 | GTTGCCCTACAGATGCCTTCC |
| TNPO3-gRNA-3 | GTCACCTGTTATTGTAACGC |
| NUP107-gRNA-1 | GTAATACTACACCAAGAAACC |
| NUP107-gRNA-2 | GAATACAGTCTGCATTAGAAG |
| NUP107-gRNA-3 | GTACTACTTTAGGAGTGGCTT |
| NUP98-gRNA-1 | GTTTGGCACAACTTCAACATT |
| NUP98-gRNA-2 | GACACCCTTTGGGGGTGGCAC |
| NUP98-gRNA-3 | GGAAATTCACAGACTAAACC |

| | |
|---|---|
| NUP214-gRNA-1 | GTTCCAAAGCCTGTTTCCAAG |
| NUP214-gRNA-2 | GTCTCCTCAGAAGTGCCCACT |
| NUP214-gRNA-3 | GTCCCCAATTCCAGCCATTAC |
| NUP62-gRNA-1 | GAACAGCGACTCTTGCTTCGG |
| NUP62-gRNA-2 | GCTCCATTCTCGATCAGCGTG |
| NUP62-gRNA-3 | GTCCAAAATTAAACCCGCTCA |
| SMARCB1-gRNA-1 | GCTAGTCGCCTCCAGAGTGAG |
| SMARCB1-gRNA-2 | GCATGCTCCACAACCATCAAC |
| SMARCB1-gRNA-3 | GTCTTCTTGTCTCGGCCCATG |
| TP53BP1-gRNA-1 | GTCCAATCCTGAACAAACAGC |
| TP53BP1-gRNA-2 | GCTGCTCAATGACCTGACTGA |
| TP53BP1-gRNA-3 | GGGGAGCAGATGGACCCTAC |
| XRCC5-gRNA-1 | GCTTTGCAGCAAGAGATGATG |
| XRCC5-gRNA-2 | GGTTCAGAGAAGATTCTTCA |
| XRCC5-gRNA-3 | GTCCTCATCCACTTTAGAGAA |
| BANF1-gRNA-1 | GCGCCAACGCCAAGCAGTCCC |
| BANF1-gRNA-2 | GAAGCAGTCCCGGGACTGCT |
| BANF1-gRNA-3 | GCCAAAAGCACCGAGACTTCG |
| EXO1-gRNA-1 | GATAGCTCTTCAAATAGCACT |
| EXO1-gRNA-2 | GAACCCGTTGATGTAATCCTC |
| EXO1-gRNA-3 | GCCTGAAACACTAAGCTACGC |
| NUP153-gRNA-1 | GCACAGGGGAAAGTGTTCCAA |
| NUP153-gRNA-2 | GAGTGAGAACGTTCAGCTTC |
| NUP153-gRNA-3 | GAAGATGTCAAGCCCTTTAG |
| PSIP1-gRNA-1 | GTCAAGATTCCAAAACCAAG |
| PSIP1-gRNA-2 | GCTTTCTCTTTCTCCCCCTTC |
| PSIP1-gRNA-3 | GGCAAACCAAATAAAAGAAA |
| XRCC6-gRNA-1 | GACCTGTATCTCGGAGATCAC |
| XRCC6-gRNA-2 | GAGGTTCGCGCCAAGGAGACC |
| XRCC6-gRNA-3 | GAATTCAAGATGAGTCATAAG |
| WRN-gRNA-1 | GACAGATGTTGCCAATAAAA |
| WRN-gRNA-2 | GTAGGAATTGAAGGAGATCAG |
| WRN-gRNA-3 | GAGCATCGTAACTATACACAA |
| PML-gRNA-1 | GGACCCTATTGACGTTGACC |
| PML-gRNA-2 | GACCGCACCCCTACGCTGACC |
| PML-gRNA-3 | GTCGGTGTACCGGCAGATTG |
| NFKB1-gRNA-1 | GAGAAGTATTTCAACCACAGA |
| NFKB1-gRNA-2 | GCCTGTTGGCAGTGCCATCTG |
| NFKB1-gRNA-3 | GCGTTTCCGTTATGTATGTGA |
| NFATC2-gRNA-1 | GCAAGTCCATACCTGAACCAC |
| NFATC2-gRNA-2 | GCGCCATGAACGCCCCCGAG |
| NFATC2-gRNA-3 | GCTCGTAAGAGCCTGACTGAC |
| cdk9-gRNA-1 | GCACCGCAAGACCGGCCAGA |
| cdk9-gRNA-2 | GACACTCTGGTACCGGCCCC |

| | |
|---|---|
| cdk9-gRNA-3 | GACTACCCTTGCAGCGGTTAT |
| RELA-gRNA-1 | GACTTACCTGCCGGGAAGATG |
| RELA-gRNA-2 | GGACAGATCAATGGCTACAC |
| RELA-gRNA-3 | GGCGCTCAGTTTCCAGAACC |
| ELL2-gRNA-1 | GCATCCAGCAAACATTCTCC |
| ELL2-gRNA-2 | GCTTACCTGGAGAATGTTTGC |
| ELL2-gRNA-3 | GACACGAGAAAGAATGACCC |
| HSP90AA1-gRNA-1 | GGTTGAGACGTTCGCCTTTC |
| HSP90AA1-gRNA-2 | GCTGCTCATCATCGTTATGTT |
| HSP90AA1-gRNA-3 | GACGATGATGAGCAGTACGCT |
| MLLT3-gRNA-1 | GACTTATTCCTGCATCTTGA |
| MLLT3-gRNA-2 | GCCAGATTCTTCTACTTTGTA |
| MLLT3-gRNA-3 | GCCTTACAAAGTAGAAGAATC |
| NFATC1-gRNA-1 | GCGGCTGCGGTCTTCGGGAG |
| NFATC1-gRNA-2 | GCGTCCACTTACAGCATTCGA |
| NFATC1-gRNA-3 | GTGGGTTGCGGGACGTGAACG |
| RICTOR-gRNA-1 | GAAAACTGGGCTTTCACTATG |
| RICTOR-gRNA-2 | GTGACAGCTTCTTTGTGATAT |
| RICTOR-gRNA-3 | GCCTTAGTAAAGTTATTCAGA |
| cdk7-gRNA-1 | GCATAGATCAGAAGCTAAAGA |
| cdk7-gRNA-2 | GATACTCACACCAATTATATT |
| cdk7-gRNA-3 | GATTTATGTCCAAAAGCATCA |
| MTOR-gRNA-1 | GCCAGCTCAGATGCCAATGAG |
| MTOR-gRNA-2 | GCTCCAGCACTATGTCACCA |
| MTOR-gRNA-3 | GAAGCACCTCTCGGAGTTCCA |
| HSF1-gRNA-1 | GCAACAGAAAGTCGTCAACA |
| HSF1-gRNA-2 | GAACTCCGTGTCGTCTCTCTC |
| HSF1-gRNA-3 | GTGAAGACATAAAGATCCGCC |
| MLLT1-gRNA-1 | GCTCCTCACCTGATTGTCCA |
| MLLT1-gRNA-2 | GGCGCCAGCCATGGACAATC |
| MLLT1-gRNA-3 | GCCCCGACTCCTCTACTTTGT |
| NFATc3-gRNA-1 | GAGCTCACTTACAGCACTCAA |
| NFATc3-gRNA-2 | GCAATAGATGGACGACCTCAG |
| NFATc3-gRNA-3 | GCTCTTTAGTATTGATTGTGC |
| PRKCD-gRNA-1 | GAAACCCTGATATATCCCAAC |
| PRKCD-gRNA-2 | GAGTTCTTACCCACGTCCTCC |
| PRKCD-gRNA-3 | GTTCCCAACGATGAACCGCCG |
| SP1-gRNA-1 | GCAGGCTCGAAGTAGCAGCAC |
| SP1-gRNA-2 | GCTACTTCGAGCCTGTGAAA |
| SP1-gRNA-3 | GAGCCCTTATTACCACCAATA |
| TBP-gRNA-1 | GACAAGGCCTTCTAACCTTAT |
| TBP-gRNA-2 | GAGAACTGAAAATCAGTGCCG |
| TBP-gRNA-3 | GAAACGCCGAATATAATCCCA |
| NFAT5-gRNA-1 | GCTACAGATGTTTCTCTAGA |

| | |
|---|---|
| NFAT5-gRNA-2 | GTGTCTATGATCTTCTCCCAA |
| NFAT5-gRNA-3 | GACAAACACTTGCAACACTAC |
| MLST8-gRNA-1 | GTCCCAGCAGGTGAATGCCT |
| MLST8-gRNA-2 | GCGCAGCATGATTGCTGCTGC |
| MLST8-gRNA-3 | GATCGATGTGGGCGGACGTGA |
| CCNT1-gRNA-1 | GCAGTCCTTCACACAGTTCCC |
| CCNT1-gRNA-2 | GCCTTCCTGATACTAGAAGTG |
| CCNT1-gRNA-3 | GATTTCCAGGGAACTGTGTGA |
| PRKCE-gRNA-1 | GAACGCATGCGGCCGAGGAAG |
| PRKCE-gRNA-2 | GATCACATACACTCTTCCTTC |
| PRKCE-gRNA-3 | GAAGCAGGGATACCAGTGTCA |
| ELL-gRNA-1 | GGAGAACGTCCGCGCCTCTG |
| ELL-gRNA-2 | GCCATGCTCTGCCGCGCCTTC |
| ELL-gRNA-3 | GGCACCGCGTCTGTTGCACC |
| AFF4-gRNA-1 | GCTTTCATACGCAGCACATTC |
| AFF4-gRNA-2 | GTCTTTCAAGGCATCAGCTTC |
| AFF4-gRNA-3 | GTTGTTCTCCAGTGCCAAAAT |
| PRKCA-gRNA-1 | GAAGTTCAAAATCCACACTTA |
| PRKCA-gRNA-2 | GAAGAAAAAGTAACAAATTCA |
| PRKCA-gRNA-3 | GTTCAGGGGGTTTGGGAAACA |
| PRKCZ-gRNA-1 | GTTCCTCACAGAGCTCCTCGA |
| PRKCZ-gRNA-2 | GCTTCGCTGTCCACCCACTTG |
| PRKCZ-gRNA-3 | GCCGAGCACCCCTGAGCAGCC |
| RPTOR-gRNA-1 | GCATGCAGAAATGTGTCAGTC |
| RPTOR-gRNA-2 | GATCAAATCCAGTGTGACGCC |
| RPTOR-gRNA-3 | GCCAGCATTCCAAGCGTGCAC |
| MATR3-gRNA-1 | GTAATTACCCAGATTTCCTCT |
| MATR3-gRNA-2 | GTAGACTGCTGCAACATGAA |
| MATR3-gRNA-3 | GAGTGGAGTCAACATATCAA |
| BCL2-gRNA-1 | GTGTGTGTGGAGAGCGTCAAC |
| BCL2-gRNA-2 | GAGAACAGGGTACGATAACC |
| BCL2-gRNA-3 | GAACTGTACGGCCCCAGCATG |
| NT-gRNA-1 | GACGGAGGCTAAGCGTCGCAA |
| NT-gRNA-2 | GCGCTTCCGCGGCCCGTTCAA |
| NT-gRNA-3 | GATCGTTTCCGCTTAACGGCG |
| NT-gRNA-4 | GTAGGCGCGCCGCTCTCTAC |
| NT-gRNA-5 | GCCATATCGGGGCGAGACATG |
| NT-gRNA-6 | GTACTAACGCCGCTCCTACAG |
| NT-gRNA-7 | GTGAGGATCATGTCGAGCGCC |
| NT-gRNA-8 | GGGCCCGCATAGGATATCGC |
| NT-gRNA-9 | GTAGACAACCGCGGAGAATGC |
| NT-gRNA-10 | GACGGGCGGCTATCGCTGACT |
| NT-gRNA-11 | GCGCGGAAATTTTACCGACGA |
