## Supplementary material for "Traitor-virus-guided discovery of novel antiviral factors": Table S2

**Supplementary Table 2. Primers used for Illumina Sequencing**

| | |
|---|---|
| NGS Fw1 | 5'-AATGATACGGCGACCACCGAGATCTACACTCTTTCCCTACACGACGCTCTTCCGATCTTAAGTAGAGTCTTGTGGAAAGGACGAAACACC-3' |
| NGS Fw2 | 5'-AATGATACGGCGACCACCGAGATCTACACTCTTTCCCTACACGACGCTCTTCCGATCTATCATGCTTATCTTGTGGAAAGGACGAAACACC-3' |
| NGS Fw3 | 5'-AATGATACGGCGACCACCGAGATCTACACTCTTTCCCTACACGACGCTCTTCCGATCTGATGCACATCTTCTTGTGGAAAGGACGAAACACC-3' |
| NGS Fw4 | 5'-AATGATACGGCGACCACCGAGATCTACACTCTTTCCCTACACGACGCTCTTCCGATCTCGATTGCTCGACTCTTGTGGAAAGGACGAAACACC-3' |
| NGS Fw5 | 5'-AATGATACGGCGACCACCGAGATCTACACTCTTTCCCTACACGACGCTCTTCCGATCTTCGATAGCAATTCTCTTGTGGAAAGGACGAAACACC-3' |
| NGS Fw6 | 5'-AATGATACGGCGACCACCGAGATCTACACTCTTTCCCTACACGACGCTCTTCCGATCTTCGATAGCAATTCTCTTGTGGAAAGGACGAAACACC-3' |
| NGS Fw7 | 5'-AATGATACGGCGACCACCGAGATCTACACTCTTTCCCTACACGACGCTCTTCCGATCTTCGATAGCAATTCTCTTGTGGAAAGGACGAAACACC-3' |
| NGS Fw8 | 5'-AATGATACGGCGACCACCGAGATCTACACTCTTTCCCTACACGACGCTCTTCCGATCTCGATCGATTTGAGCCTTCTTGTGGAAAGGACGAAACACC-3' |
| NGS Fw9 | 5'-AATGATACGGCGACCACCGAGATCTACACTCTTTCCCTACACGACGCTCTTCCGATCTACGATCGATACACGATCTCTTGTGGAAAGGACGAAACACC-3' |
| NGS Fw10 | 5'-AATGATACGGCGACCACCGAGATCTACACTCTTTCCCTACACGACGCTCTTCCGATCTTACGATCGATGGTCCAGATCTTGTGGAAAGGACGAAACACC-3' |
| NGS Rv1 | 5'-CAAGCAGAAGACGGCATACGAGATTCGCCTTGGTGACTGGAGTTCAGACGTGTGCTCTTCCGATCTGCCCTTATTTTAACTTGCTATTTCTAGCTCTAAAAC-3' |
| NGS Rv2 | 5'-CAAGCAGAAGACGGCATACGAGATATAGCGTCGTGACTGGAGTTCAGACGTGTGCTCTTCCGATCTGCCCTTATTTTAACTTGCTATTTCTAGCTCTAAAAC-3' |
| NGS Rv3 | 5'-CAAGCAGAAGACGGCATACGAGATGAAGAAGTGTGACTGGAGTTCAGACGTGTGCTCTTCCGATCTGCCCTTATTTTAACTTGCTATTTCTAGCTCTAAAAC-3' |
| NGS Rv4 | 5'-CAAGCAGAAGACGGCATACGAGATATTCTAGGGTGACTGGAGTTCAGACGTGTGCTCTTCCGATCTGCCCTTATTTTAACTTGCTATTTCTAGCTCTAAAAC-3' |
| NGS Rv5 | 5'-CAAGCAGAAGACGGCATACGAGATCGTTACCAGTGACTGGAGTTCAGACGTGTGCTCTTCCGATCTGCCCTTATTTTAACTTGCTATTTCTAGCTCTAAAAC-3' |

| | |
|---|---|
| NGS Rv6 | 5’-CAAGCAGAAGACGGCATACGAGATGTCTGATGGTGACTGGAGTTCAGACGTGTGCTCTTCCGATCTGCCCTTATTTTAACTTGCTATTTCTAGCTCTAAAAC-3’ |
| NGS Rv7 | 5’-CAAGCAGAAGACGGCATACGAGATTTACGCACGTGACTGGAGTTCAGACGTGTGCTCTTCCGATCTGCCCTTATTTTAACTTGCTATTTCTAGCTCTAAAAC-3’ |
| NGS Rv8 | 5’-CAAGCAGAAGACGGCATACGAGATTTGAATAGGTGACTGGAGTTCAGACGTGTGCTCTTCCGATCTGCCCTTATTTTAACTTGCTATTTCTAGCTCTAAAAC-3’ |
